# DNA replication fork speed Acts as a Pacer in Cortical Neurogenesis

**DOI:** 10.1101/2024.11.05.622174

**Authors:** Jianhong Wang, Yifan Kong, Xuezhuang Li, Kun Xiang, Yulian Tan, Lei Shi

## Abstract

DNA replication fork speed, which controls the rate of genome duplication, has emerged as a key regulator of cellular plasticity. However, its role in neurogenesis remains unexplored. Mini-chromosome maintenance complex (MCMs)-binding protein (MCMBP) functions as a chaperone for newly synthesized MCMs, increasing chromatin coverage to restrain fork speed. We demonstrate that selective deletion of MCMBP in neural progenitor radial glial cells (RGCs) accelerates fork speed, triggering widespread apoptosis, DNA damage, and micronuclei formation, ultimately activating p53 and causing microcephaly. Unexpectedly, concurrent deletion of *Trp53* and *Mcmbp* further increased fork speed, leading to extensive RGC detachment from the ventricular zone and acquisition of outer-RGC characteristics. Mechanistically, we found that the MCM3 subunit coordinates DNA and centrosome duplication, thereby mediating RGC attachment. Behavioral analysis revealed that disruption of fork speed results in anxiety-like behavior in mice. These findings unveil a previously unrecognized role for replication fork speed in neurogenesis.

## Introduction

The neocortex, a hallmark of mammalian evolution, orchestrates vital functions including sensation, emotion, sleep, learning, and memory ^1–3^. This complex structure develops through precisely coordinated neurogenic events, with genome integrity maintenance being crucial for proper cortical function. DNA replication is fundamental to cortical development ^4,5^, characterized by neural progenitor cells (NPCs) exhibiting progressively longer S-phase durations as neurogenesis proceeds ^4^. This temporal pattern indicates tight regulation of DNA fork progression rates during neurogenesis, although the underlying regulatory mechanisms and consequences of their disruption remain poorly understood.

Cortical development initiates with neural epithelial stem cells, which undergo active self-amplification to expand the neural progenitor cell population before transitioning into the primary NPCs known as apical radial glial cells (RGCs) ^6,7^. Within the ventricular zone (VZ), apical RGCs exhibit distinct polarity, featuring a short apical process anchored to the ventricular surface and a long basal process that extends to the cortical surface ^6^. During early neurogenesis, RGCs primarily undergo symmetric divisions to generate two identical RGCs ^6–8^. As neurogenesis progresses, RGCs gradually transition to asymmetric divisions, either directly generating neurons or producing intermediate progenitor cells (IPCs) in the sub-ventricular zone (SVZ) that subsequently generate neurons indirectly ^6–8^.

DNA replication is a highly regulated process that initiates during G1 phase with the assembly of pre-replicative complexes (pre-RCs) at replication origins ^9^. The replicative complex (RC), evolutionarily conserved from yeast to humans, consists of MCM2-7 proteins arranged in a hexameric ring ^10,11^. During S phase, mini-chromosome maintenance proteins (MCMs) form an excess of DNA replication complexes, with only a fraction becoming active CDC45–MCM–GINS (CMG) helicases that bind to replication origins and initiate DNA replication ^10,12^. Proliferating cells maintain high MCM expression levels to ensure robust and faithful DNA replication ^13^.

Two functionally distinct pools of MCMs are inherited by daughter cells ^13,14^: (1) Parent MCMs, which participated in DNA replication in mother cells, and (2) Nascent MCMs, newly formed in mother cells, which regulate CMG helicase movement by increasing chromatin coverage, thereby reducing DNA replication errors ^13,14^. MCM2 enters the nucleus independently and subsequently interacts with MCM3-7 to form complete MCM complexes ^15,16^. MCM binding protein (MCMBP) serves as a dedicated chaperone essential for MCM3-7 organization, facilitating both assembly and unloading of newly synthesized MCM3-7 subcomplexes with MCM2 to form nascent MCMs ^10,17^ (**Fig. 1a**). In the absence of MCMBP, MCM3-7 subunits undergo cytoplasmic degradation, resulting in inefficient nuclear import ^14,18^.

**Fig.1.**
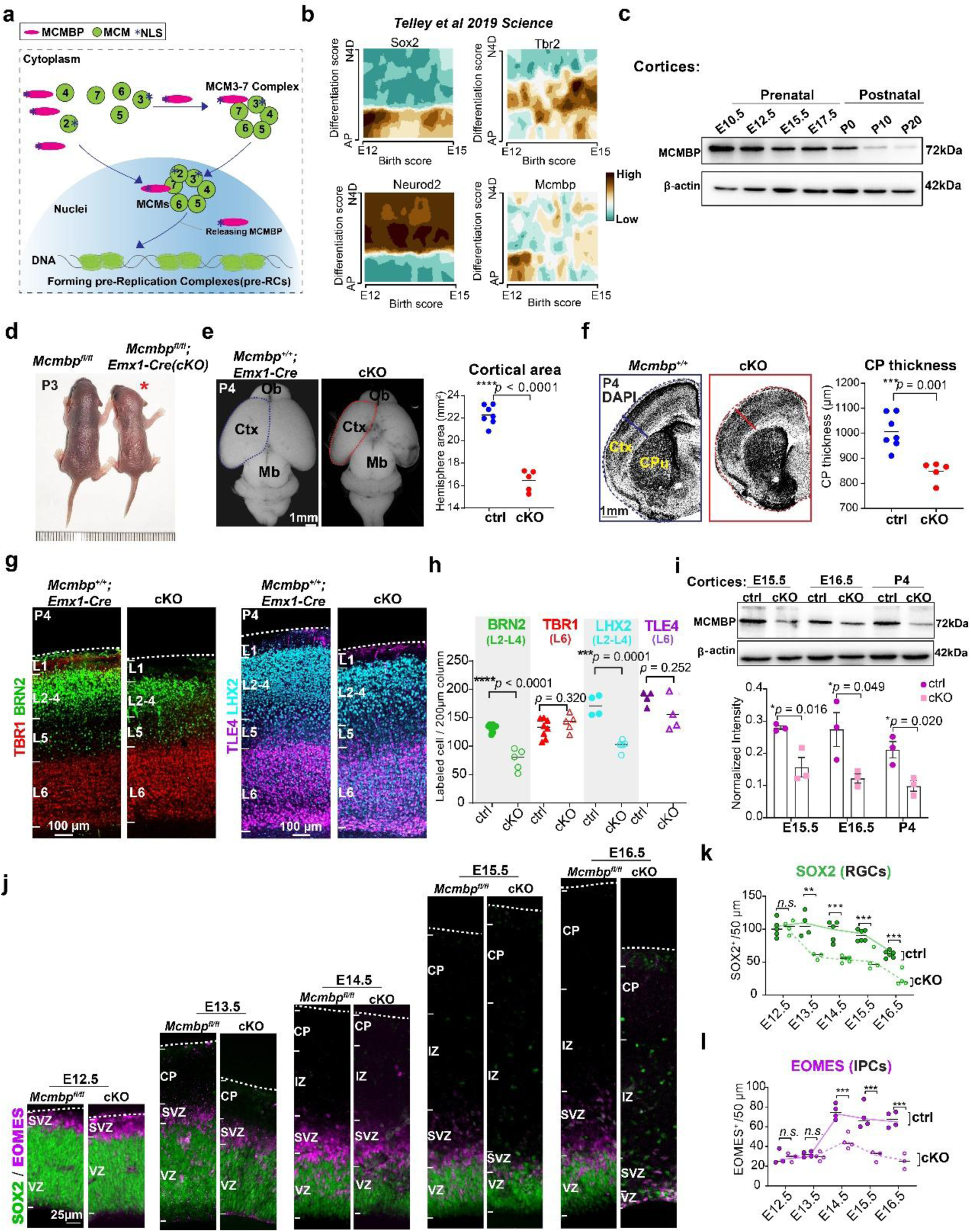
|*Mcmbp* deletion in neural progenitors leads to microcephaly. **a,** Schematic map showing MCMBP-mediated assembly, and export MCM3-7 into nuclear which forms nascent MCMs with MCM2 as pre-replication complexes, and regulates DNA replication fork speed. NLS means nuclei localization signal. **b,** Spatial-temporal expression of *Mcmbp* from apical to basal position, and from E12.5 to E15.5. **c,** Western blot analysis showing MCMBP expression pattern during prenatal and postnatal stages of cortical development. **d,** Representative images of cKO mouse and littermate control at P3. The red star indicates cKO mouse. **e,** (Left panel) Dorsal view of *Mcmbp^+/+^; Emx1-Cre* and *Mcmbp^fl/fl^; Emx1-Cre* (cKO) P4 brain. (Right panel) The cortical area was significantly reduced in cKO compared to littermate control (ctrl). (mean, two-tailed unpaired t-test, ctrl: n=7, cKO: n=5). **f,** (Left panel) DAPI staining coronal sections of *Mcmbp^+/+^*and cKO P4 brain. (Right panel) Cortical plate thickness was significantly reduced in cKO compared to littermate control (ctrl). (mean, two-tailed unpaired t-test, ctrl: n=7, cKO: n=5). **g,** Immunostaining of layer markers BRN2, TBR1, LHX2 and TLE4 in in P4 brain of *Mcmbp^+/+^; Emx1-Cre* and cKO. **h,** Upper layer neurons were significantly reduced in cKO compared to the littermate controls (ctrl). (mean, two-tailed unpaired t-test, BRN2, TBR1, ctrl: n=8, cKO: n=5, LHX2, TLE4, ctrl: n=4, cKO: n=4). **i,** Western blot analysis showing downregulation of MCMBP expression in the E15.5, E16.5 and P4 cortex. (mean, two-tailed unpaired t-test, ctrl: n=3, cKO: n=3). **j,** Immunostaining of apical progenitor marker SOX2 and intermediate progenitor marker EOMES in *Mcmbp^+/+^; Emx1-Cre* and cKO from E12.5 to E16.5. **k,** SOX2+ cell number analysis indicated that there was no difference between ctrl and cKO at E12.5. However, since E13.5, SOX2+ cells were significantly reduced and continued until E16.5. (mean, two-tailed unpaired t-test, E12.5, ctrl: n=5, cKO: n=4, E13.5, ctrl: n=4, cKO: n=3, E14.5, ctrl: n=5, cKO: n=5, E15.5, ctrl: n=6, cKO: n=4, E16.5, ctrl: n=6, cKO: n=4). **l,** EOMES+ cell number analysis indicated that there was no difference between ctrl and cKO at E12.5 and E13.5. However, EOMES+ cells were significantly reduced from E14.5 to E16.5. (mean, two-tailed unpaired t-test, E12.5, ctrl: n=3, cKO: n=3, E13.5, ctrl: n=4, cKO: n=4, E14.5, ctrl: n=4, cKO: n=4, E15.5, ctrl: n=4, cKO: n=3, E16.5, ctrl: n=4, cKO: n=3).

Based on these findings, we hypothesized that DNA fork speed could be manipulated during cortical development through conditional deletion of *Mcmbp* in the mouse embryonic brain, and we investigated its consequences *in vivo*. Our results demonstrate that *Mcmbp* deletion increases DNA fork speed in RGCs, triggering massive cell apoptosis, DNA damage, and robust p53 activation during later stages of neurogenesis. Surprisingly, simultaneous deletion of both *Trp53* and *Mcmbp* resulted in even faster DNA fork speeds than *Mcmbp* deletion alone, causing RGCs to gradually detach from the VZ surface and acquire features characteristic of outer RGCs. We further examined whether increased fork speed during embryonic brain development influenced adult mouse behavior. Collectively, our findings unveil a previously unrecognized role for DNA fork speed in cortical development and demonstrate its lasting impact on adult mouse behavior.

## Results

### *MCMBP* Expression Is Essential for Cortical Development

Initially, we analyzed spatial and temporal mouse brain transcriptome data ^19^ to examine *Mcmbp* expression patterns. We found that *Mcmbp* expression peaked at E12 and gradually decreased over the following 48 hours of cortical development through E15 (**Fig. 1b**). To validate these findings, we performed Western blot (WB) analysis on cortical tissue from four embryonic stages (E10.5, E12.5, E15.5, and E17.5) and three postnatal time points (postnatal days 0, 10, and 20) (**Fig. 1c**). Consistent with the RNA expression data, MCMBP protein levels progressively decreased during brain development (**Fig. 1c**), suggesting a regulatory role in cortical neurogenesis.

Re-analysis of publicly available single-cell transcriptome data from E14.5 mouse cortex ^20^ revealed significant enrichment of *Mcmbp* expression in neural progenitor cells (**Extended Data Fig. 1a**, Cluster 11, *p-adj* = 3.21E-23). These results suggest that the *Mcmbp* expression is highest in neural progenitor cells and exhibits a gradual decrease pattern during cortical development, and thus MCMBP can be manipulated to investigate the role of DNA fork speed in RGCs.

To investigate MCMBP function, we generated conditional knockout (cKO) mice using *Emx1-Cre* ^21^, which mediates Cre recombination in cortical RGCs beginning at E10.5 (**Extended Data Fig.1b, c**). While *Mcmbp^fl/fl^; Emx1-Cre* cKO mice were viable at birth, by postnatal day 3 (P3) they exhibited developmental deficits characterized by red skin and reduced body size (**Fig. 1d**). Analysis at two postnatal brain maturation stages (P4 and P8) revealed that cKO mice exhibited reduced cortical area (26.09% and 32.56% decrease, respectively; **Fig.1e** and **Extended Data Fig.1d**) and decreased cortical plate thickness (15.71% and 21.90% reduction, respectively; **Fig.1f** and **Extended Data Fig.1e**) compared to controls.

To examine the impact of *Mcmbp* deletion on different neuronal subtypes, we analyzed cortical layer markers at P4 and P8. Expression of deep layer neuronal markers TBR1 ^22^ and TLE4 ^23^ in layer (L) 6 remained unchanged in both number and laminar location (**Fig. 1g**, **h** and **Extended Data Fig.1f, g**). In contrast, upper layer (L2-4) neuronal markers showed significant reductions: BRN2 ^24^ and LHX2 ^25,26^ decreased by 41.99% and 41.11% respectively at P4 (**Fig. 1g**, **h**), with further reductions of 55.26% and 63.10% at P8 (**Extended Data Fig.1f, g**). WB analysis confirmed progressive reduction of MCMBP expression in cKO cortices compared to controls: ∼43.98% at E15.5, ∼55.29% at E16.5, and ∼53.34% at P4 (**Fig.1i**).

During sequential neurogenesis, RGCs generate deep layer neurons before producing upper layer neurons ^6^. To track this process, we quantified RGC numbers from E12.5 to E16.5 (**Fig.1j**). In cKO mice, SOX2-expressing RGCs first showed significant numerical reduction at E13.5 (**Fig.1k**). Similarly, EOMES-expressing IPCs exhibited significant reduction beginning at E14.5 (**Fig.1l**). By E16.5, when upper layer neurons are typically generated, both apical RGCs and IPCs were largely depleted (**Fig.1k, l**). These results demonstrate that *Mcmbp* deletion leads to progressive loss of neural progenitor cells. Notably, at E15.5 and E16.5, we detected SOX2+ cells outside the VZ in cKO mice, suggesting RGC delamination following *Mcmbp* deletion (**Fig.1j**).

In summary, while cortical layer organization remained intact following *Mcmbp* deletion, upper layer neuron numbers were significantly reduced due to progressive NPC loss during neurogenesis.

### *Mcmbp* deletion Induces Rapid Apoptosis and Cell Cycle Exit

To determine whether apoptosis contributed to NPC loss, we analyzed activated caspase-3 levels in the cortices. At E15.5, we observed robust cleaved caspase-3 (CC3) labeling in the VZ, SVZ, and intermediate zone (IZ) of cKO cortex, while only rare instances were detected in the cortical plate (**Extended Data Fig.2a**). DAPI staining revealed numerous pyknotic nuclei—a hallmark of apoptosis characterized by chromatin condensation—which were rarely observed in controls (**Extended Data Fig.2b**). Consistent with previous reports^27^, these pyknotic nuclei exhibited a distinctive clustering pattern (**Extended Data Fig.2b**, red square). Quantification showed that pyknotic nuclei were scarce at E12.5, progressively increased from E13.5 to E15.5, and then decreased dramatically at E16.5 (**Extended Data Fig.2c**). Temporal analysis of apoptosis revealed an unexpected pattern: apoptotic cells were rarely detected in the cortical zone at E12.5, within two days of *Emx1-Cre* onset (**Extended Data Fig.2d**). During later division stages, apoptosis emerged at E13.5 and increased gradually until E15.5, coinciding with peak neurogenesis. Remarkably, CC3 levels decreased substantially by E16.5, just one day after peak expression (**Extended Data Fig.2d**). These findings further demonstrate that *Mcmbp* expression is tightly regulated during neurogenesis, and its disruption triggers massive cellular apoptosis.

To determine the temporal dynamics of apoptosis, we tracked NPC fate using dual-pulse labeling with thymidine analogs 5-ethynyl-2′-deoxyuridine (EdU) and 5-bromo- 2′-deoxyuridine (BrdU), which incorporate into DNA during S phase. We administered BrdU 4 hours before analysis at E15.5 to label NPCs recently entering S phase. While BrdU-labeled NPCs were detected in both VZ and SVZ of cKO and littermate controls, fewer BrdU+ cells were observed in the SVZ of cKO mice (**Extended Data Fig.2e**). In cKO mice, BrdU+ cells significantly colocalized with CC3 in both VZ and SVZ (**Extended Data Fig.2e**, bottom panel), suggesting rapid onset of apoptosis following *Mcmbp* deletion. For long-term tracking of S-phase cells, we administered EdU 24 hours before analysis at E15.5. While only 6.67% of BrdU+ cells showed apoptosis (**Extended Data Fig.2e**), approximately 71.33% of EdU+ NPCs became apoptotic within 24 hours in cKO mice (**Extended Data Fig.2f**). Unexpectedly, we detected apoptotic signals in post-mitotic neurons within the cortical plate of cKO mice (**Extended Data Fig.2f**, white arrow), suggesting that *Mcmbp* deletion may also promote premature RGC differentiation.

To examine cell cycle progression, we used phospho-histone H3 (pHH3) staining to mark M-phase cells. Compared to controls, cKO mice showed fewer M-phase RGCs in both luminal-VZ and outside-VZ regions (**Extended Data Fig.3a**). We then combined Ki67 staining (marking all cycling cells) with 4-hour BrdU pulse labeling (marking S-phase cells). The proportion of BrdU+Ki67-/BrdU+ cells, representing cells exiting the cell cycle, reached 15.33% in cKO RGCs (**Extended Data Fig.3b**). After 24-hour EdU pulse labeling, an even higher percentage (∼54.01%) of RGCs had exited the cell cycle (**Extended Data Fig.3c**) but failed to properly differentiate into post-mitotic neurons. These findings indicate that cKO RGCs fail to complete S-phase and prematurely exit the cell cycle. Consistent with incomplete S-phase progression, we observed fewer neurons in the IZ, the primary location of migrating neurons (**Extended Data Fig.2f**). Together, these data demonstrated that following *Mcmbp* deletion, RGCs predominantly failed to complete the cell cycle for neuronal differentiation and underwent rapid apoptosis within 24 hours.

### *Mcmbp* Deletion Induces Severe DNA Damage and p53 Activation

To elucidate the molecular mechanisms underlying *Mcmbp* deletion-induced apoptosis, we performed transcriptome analysis on dissected cortices (control n=6 vs cKO n=6) at E15.5 (**Fig. 2a**). Differential expression analysis identified 279 upregulated and 102 downregulated genes in cKO samples (**Fig.2a** and **Supplementary Table 1**). Gene ontology (GO) analysis revealed that downregulated genes were significantly enriched in forebrain development-related terms, while upregulated genes showed enrichment in immune response pathways (**Fig.2b, c** and **Supplementary Table 2**). Notably, pathway analysis highlighted an overrepresentation of p53-mediated signaling pathways (**Fig.2d**, 4/10, highlighted in pink color), suggesting that p53 upregulation triggered the apoptotic cascade. Among the top regulated genes, we observed increased expression of *Eda2r*, a known mediator of p53-dependent cell death^28^ in cKO samples (**Fig. 2e**) - a finding subsequently validated by real-time PCR analysis (**Fig.2e**, right panel). Immunostaining for stabilized p53 revealed a marked increase in p53-positive cells in E15.5 cKO tissue, with particular enrichment in the VZ/SVZ regions (**Fig.2f, g**, cyan arrow).

**Fig.2.**
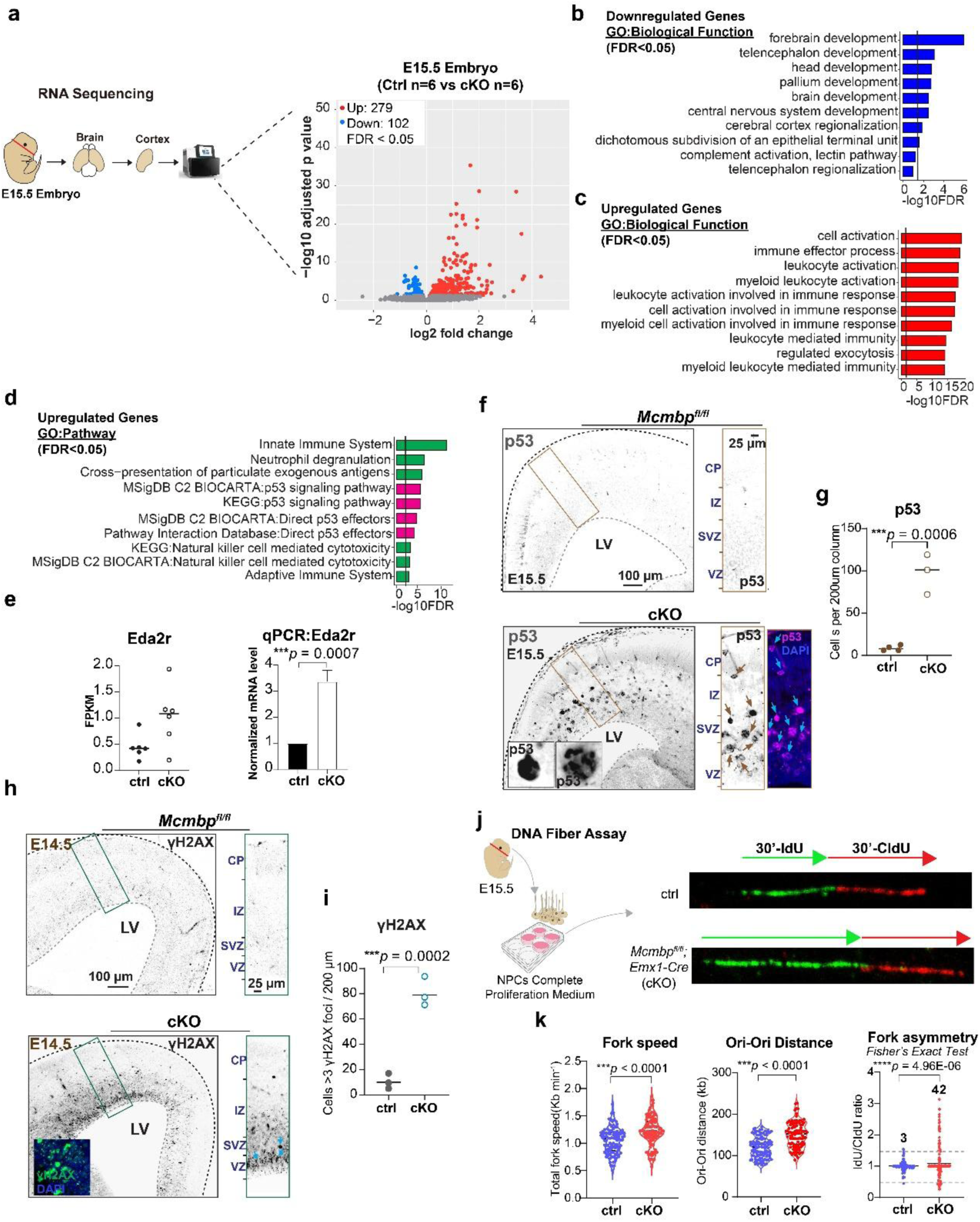
*| Mcmbp* deletion led to DNA damage and p53 activation. **a,** Schematic map of RNA-seq process at E15.5 and volcano plot of differentially expressed genes (DEGs) between cKO (n=6) and control (n=6). Down-regulated genes (FDR<0.05) are indicated in blue and up-regulated genes (FDR<0.05) are indicated in red. **b-c,** GO biological function analysis of downregulated genes and upregulated genes in cKO **d,** GO pathway analysis of upregulated genes in cKO. **e,** Q-PCR results validated Eda2r high expression pattern in cKO. (mean, two-tailed unpaired t-test, ctrl: n=3, cKO: n=3). **f-g,** Immunostaining and quantification of p53 at E15.5, revealed p53+ cells were significantly increased in cKO. (mean, two-tailed unpaired t-test, ctrl: n=4, cKO: n=3). **h-i,** Immunostaining and quantification of γH2AX+ at E14.5, revealed γH2AX+ cells were significantly increased in cKO. (mean, two-tailed unpaired t-test, ctrl: n=3, cKO: n=3). **j-k,** DNA fiber analysis between control and cKO at E15.5 indicated that DNA fork speed (mean, two-tailed unpaired t-test, ctrl: n=100, cKO: n=100), ori-ori distance (mean, two-tailed unpaired t-test, ctrl: n=114, cKO: n=104), and fork asymmetry (Fisher’s Exact Test, ctrl: n=100, cKO: n=150) were significantly increased in cKO.

Disruption of DNA fork progression may increase susceptibility to DNA damage, potentially triggering RGC apoptosis following *Mcmbp* deletion. To evaluate DNA damage during neurogenesis, we analyzed the DNA damage marker γH2AX through immunostaining and quantification at three developmental timepoints: E14.5, E15.5, and E16.5. The analysis revealed a significant increase in γH2AX-positive cells (>3 foci) in cKO compared to controls, with γH2AX signals predominantly concentrated in the VZ/SVZ regions where NPCs reside (**Fig.2h** and **Extended Data Fig.4a, b**). Notably, DNA damage peaked at E14.5, one day before maximum apoptosis in cKO, with an average of 80 cells/200μm displaying >3 γH2AX foci, compared to lower levels at E15.5 and E16.5 (40 cells/200μm >3 foci for both timepoints) (**Fig. 2i** and **Extended Data Fig. 4a, b**).

To assess DNA replication fork speed in cKO at E15.5, we performed DNA fiber analysis. Primary cortical NPCs were isolated from E15.5 embryonic cortex and cultured in complete proliferation medium for 1-2 days. Active replication forks were labeled using sequential incorporation of two distinct halogenated nucleotides: IdU (5- Iodo-2′-deoxyuridine) and CldU (5-Chloro-2′-deoxyuridine) into nascent DNA strands (**Fig. 2j**). Comparative analysis demonstrated significantly increased DNA fork speed following *Mcmbp* deletion (**Fig. 2k**, control mean = 1.048 kb/min versus cKO mean = 1.217 kb/min). Origin firing assessment, measured by inter-origin distances, revealed an increased average distance in cKO (149.8 kb) compared to controls (120.7 kb) (**Fig. 2k**, middle panel). This finding aligns with reduced *Mcmbp* expression and decreased replication origin numbers, ultimately resulting in significantly enhanced fork asymmetry in cKO (**Fig. 2k**, right panel).

Collectively, these findings demonstrate that *Mcmbp* deletion disrupts DNA fork speed regulation in RGCs, leading to increased fork asymmetry, extensive DNA damage response, and micronuclei formation during S phase in daughter cells, ultimately triggering widespread p53-dependent apoptosis during cortical development.

### *Trp53/Mcmbp* Co-deletion Substantially Rescues the Microcephaly Phenotype

To further elucidate p53’s role in mediating apoptotic response and the impact of accelerated fork speed during neurogenesis, we generated *Mcmbp/Trp53* conditional double mutants (*Mcmbp^fl/fl^; Trp53^fl/fl^; Emx1-Cre*, dKO). Remarkably, dKO mice at P3 exhibited normalized body size and skin coloration comparable to controls (**Fig. 3a**). By P8, both brain size and cortical thickness showed substantial recovery (**Fig. 3b, c**). Analysis of cortical layering revealed that upper layer neurons (L2-4) were largely restored following p53 deletion in cKO, while deep layer neurons (L5-L6) remained unchanged across all groups - controls, cKO, and dKO (**Fig. 3d-g**). Notably, we observed aberrant distribution of LHX2-positive upper layer neurons throughout the cortical plate, with some cells mislocalized to deep layers in dKO (**Fig. 3f**, white star), suggesting impaired neuronal migration.

**Fig. 3.**
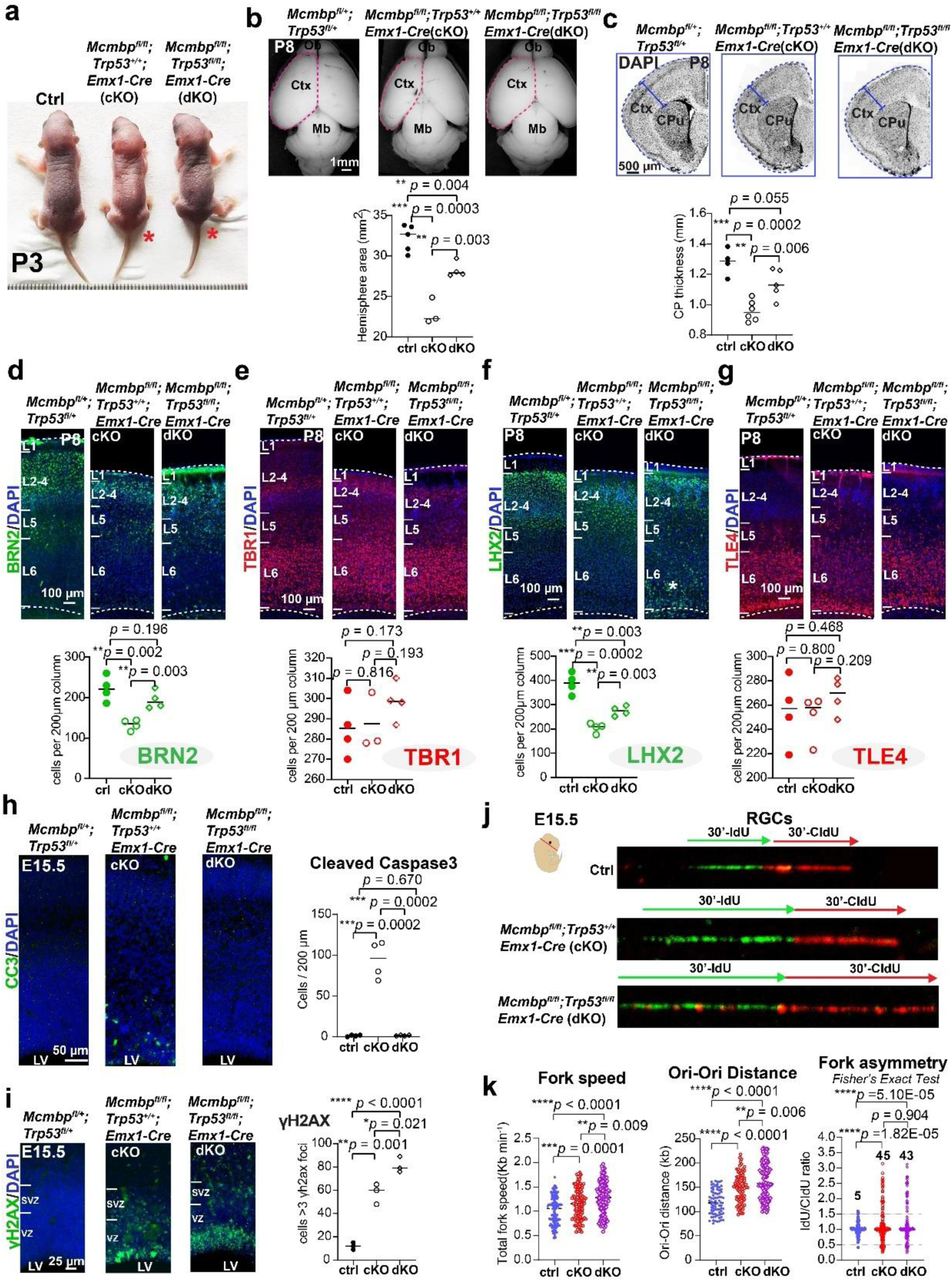
| *Trp53/Mcmbp* co-deletion partially rescued microcephaly. **a,** Representative images of *Mcmbp^fl/fl^; Emx1*(cKO), *Mcmbp^fl/fl^; Trp53^fl/fl^; Emx1*(dKO) mice, and littermate controls at P3. Red star indicates *Mcmbp* cKO and dKO. **b,** (Top panel) Dorsal view of control, cKO, and dKO brain at P4. (Bottom panel) Cortical area was partially rescued in dKO compared to littermate control (ctrl) and cKO. (mean, two-tailed unpaired t-test, ctrl: n=5, cKO: n=3, dKO: n=4). **c,** (Top panel) DAPI staining coronal sections of *Mcmbp ^fl/+^; Trp53 ^fl/+^,* cKO and dKO P4 brain. (Bottom panel) Cortical plate thickness was rescued in dKO compared to littermate control and cKO. (mean, two-tailed unpaired t-test, ctrl: n=4, cKO: n=6, dKO: n=5). **d-g,** (Top panel) Immunostaining of layer markers BRN2(d), TBR1(e), LHX2(f), TLE4(g) in *Mcmbp ^fl/+^; Trp53 ^fl/+^*, cKO, and dKO P8 brain. (Bottom panel) Upper layer neurons BRN2, but not LHX2, were rescued in dKO, deeper layer neurons showed no differences compared to the control and cKO. (mean, two-tailed unpaired t-test, BRN2, LHX2, TLE4, ctrl: n=4, cKO: n=4, dKO: n=4, TBR1, ctrl: n=4, cKO: n=3, dKO: n=4). **h,** Immunostaining and quantification results of CC3+ indicated that apoptotic cells were completely eliminated in dKO. (mean, two-tailed unpaired t-test, ctrl: n=4, cKO: n=4, dKO: n=4). **i,** Immunostaining and quantification results of γH2AX+ indicated that DNA-damaged cells were significantly increased in dKO compared to control and cKO. (mean, two-tailed unpaired t-test, ctrl: n=3, cKO: n=3, dKO: n=3). **j-k,** DNA fiber analysis among control, cKO, and dKO at E15.5 indicated that DNA fork speed (mean, two-tailed unpaired t-test, ctrl: n=130, cKO: n=130, dKO: n=130) and the origin-origin distance (mean, two-tailed unpaired t-test, ctrl: n=78, cKO: n=81, dKO: n=116) were significantly increased in dKO compared to both control and cKO. The dKO fork asymmetry is comparable to cKO. (Fisher’s Exact Test, ctrl: n=100, cKO: n=150, dKO: n=150).

To determine whether the phenotypic rescue stemmed from prevented apoptosis in p53’s absence, we quantified CC3-positive cells. The dKO showed comparable numbers to controls (**Fig. 3h**), confirming that *Mcmbp* deletion-induced apoptosis is p53-dependent. We then assessed the persistence of DNA damage in dKO. Immunostaining revealed significantly higher numbers of γH2AX-positive cells (>3 foci) in dKO compared to cKO (**Fig. 3i**, dKO 81.33%, cKO 57.67%, control 12%), indicating that *Mcmbp* deletion-induced DNA damage persists despite p53 deletion. However, the number of phospho-(p) KAP1 positive cells - an ATM substrate and marker of heterochromatic double-strand breaks (DSBs) ^29^ - returned to normal levels in dKO (**Extended Data Fig. 5**), suggesting p53’s involvement in DSB formation during DNA replication.

To further characterize the impact of p53 deletion, we examined fork speed in dKO using fiber assay experiments (**Fig. 3j**). Analysis revealed that RGCs in dKO exhibited significantly higher fork speeds compared to cKO (**Fig. 3k**, left panel). Concurrent with increased fork speed, the inter-origin distance was markedly elevated in dKO, averaging 163.7 kb (**Fig. 3k**, middle panel). Regarding fork asymmetry, both cKO and dKO showed significant increases compared to controls, though no significant difference was observed between cKO and dKO (**Fig. 3k**, right panel).

### *Trp53/Mcmbp* Co-deletion Induces Extensive RGC Detachment from the Ventricular Zone

We next investigated whether neural progenitors were rescued in p53’s absence. While the total number of SOX2-positive apical RGCs increased in dKO (**Fig. 4a, b**), we unexpectedly observed extensive ectopic localization of SOX2-positive cells outside the ventricular zone (**Fig. 4a**, bottom panel). Distribution analysis revealed that these displaced SOX2-positive progenitors predominantly accumulated in the SVZ and IZ - regions typically enriched with intermediate progenitor cells and migrating neurons, respectively (**Fig. 4c**).

**Fig. 4.**
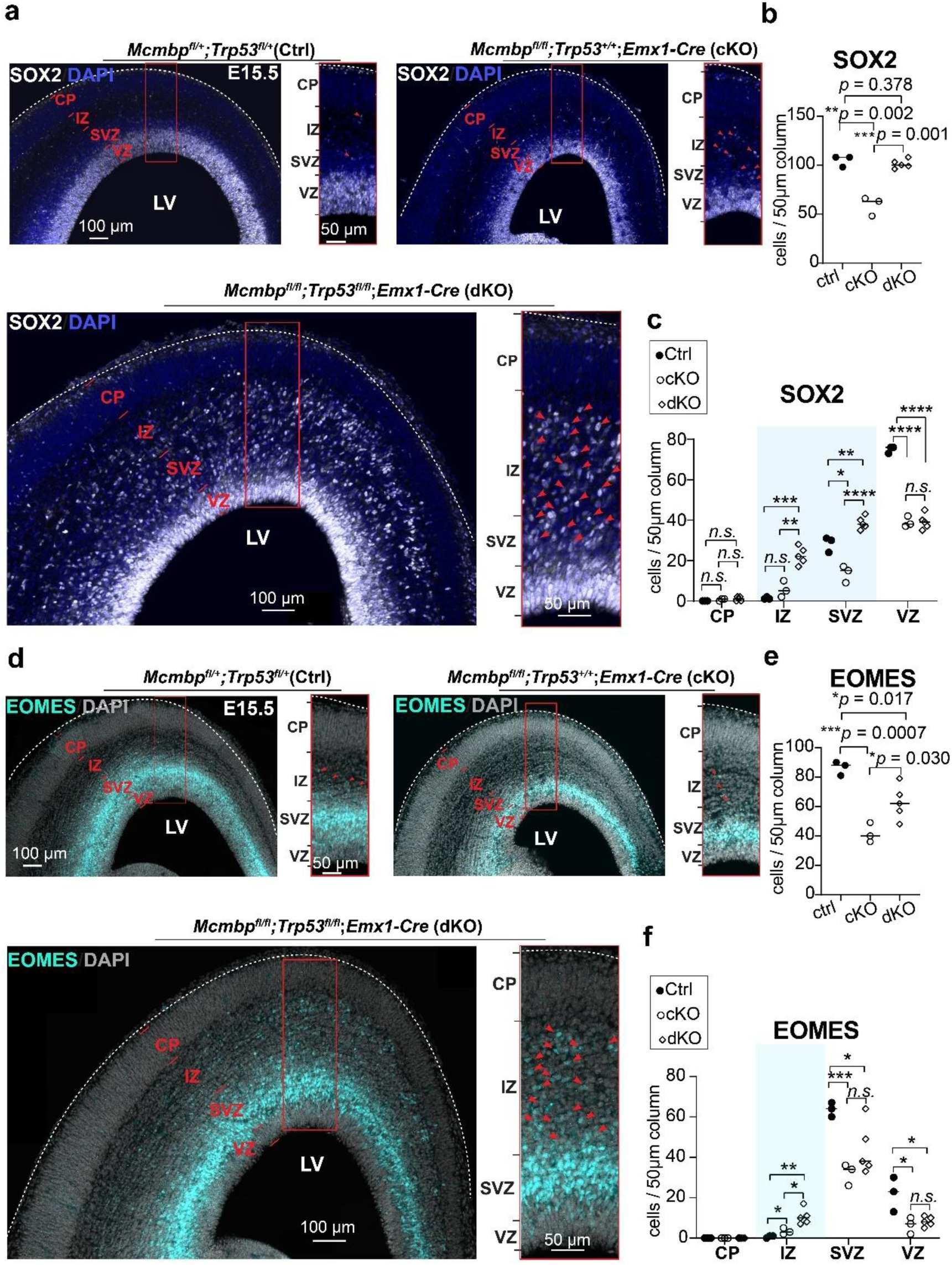
|*Trp53/Mcmbp* co-deletion leads to delocalization of RGCs and IPCs. **a,** Immunostaining of E15.5 cortex revealed that massive SOX2+ cells (white color) were displaced in extra-VZ in dKO. Red arrows indicate displaced SOX2+ cells. **b,** Quantification of the SOX2+ cells revealed that dKO SOX2+ cells were rescued to levels comparable to controls. (mean, two-tailed unpaired t-test, ctrl: n=3, cKO: n=3, dKO: n=5). **c,** Quantification of SOX2+ cell distribution revealed that displaced SOX2+ cells in dKO were mainly in SVZ and IZ, highlighted in blue. (mean, two-tailed unpaired t-test, ctrl: n=3, cKO: n=3, dKO: n=5). **d,** Immunostaining of E15.5 cortex revealed the EOMES+ (cyan color) cells were displaced in extra-SVZ in dKO. Red arrow indicates displaced EOMES+ cells. **e,** Quantification of the EOMES+ cells revealed that dKO EOMES+ cells were higher than cKO, but less than controls. (mean, two-tailed unpaired t-test, ctrl: n=3, cKO: n=3, dKO: n=5). **f,** Quantification of EOMES+ cell distribution revealed that dKO EOMES+ cells were displaced in IZ highlighted with cyan color. (mean, two-tailed unpaired t-test, ctrl: n=3, cKO: n=3, dKO: n=5).

Paralleling the apical RGC findings, EOMES-positive intermediate progenitors (IPs) showed significantly higher numbers in dKO compared to cKO (**Fig. 4d**, **e**), though remaining slightly below control levels (**Fig. 4e**). Notably, the majority of these increased IPs exhibited ectopic localization beyond the SVZ, with particular enrichment in the IZ (**Fig. 4d**, bottom panel, and **4f**), suggesting their possible differentiation from displaced SOX2-positive cells.

Temporal analysis of RGCs across developmental stages (E13.5, E14.5, E15.5, and E17.5) revealed progressive detachment of both SOX2-positive RGCs and EOMES- positive IPs from the ventricular surface (**Extended Data Fig. 6**). Remarkably, even at E17.5, near neurogenesis completion, substantial numbers of displaced SOX2-positive RGCs remained detectable within the IZ (**Extended Data Fig. 6**).

### Detached Progenitors Display Shortened S-phase Duration and Enhanced Proliferative Capacity

To characterize transcriptional changes following *Trp53* and *Mcmbp* co-deletion, we performed RNA-seq analysis of *Mcmbp/Trp53* dKO cortex at E15.5. This analysis identified 230 upregulated and 56 downregulated genes (**Fig. 5a** and **Supplementary Table 3**). Comparison with cKO-derived differentially expressed genes (DEGs) revealed minimal overlap, with only 19 upregulated and 4 downregulated genes shared between conditions (**Fig. 5b** and **Supplementary Table 4**), indicating that *Trp53* deletion largely restores normal gene expression patterns. Consistent with these findings, *Trp53* expression was significantly reduced in dKO, while Eda2r levels remained comparable to controls (**Fig. 5c**). Gene ontology analysis revealed enrichment of upregulated genes in pathways related to cell cycle progression, cell division, and chromosome segregation (**Fig. 5d**, top panel and **Supplementary Table 5**). Molecular function analysis further demonstrated that upregulated genes primarily participate in DNA replication and DNA helicase activity **(Fig.5d**, bottom panel and **Supplementary Table 5**).

**Fig. 5.**
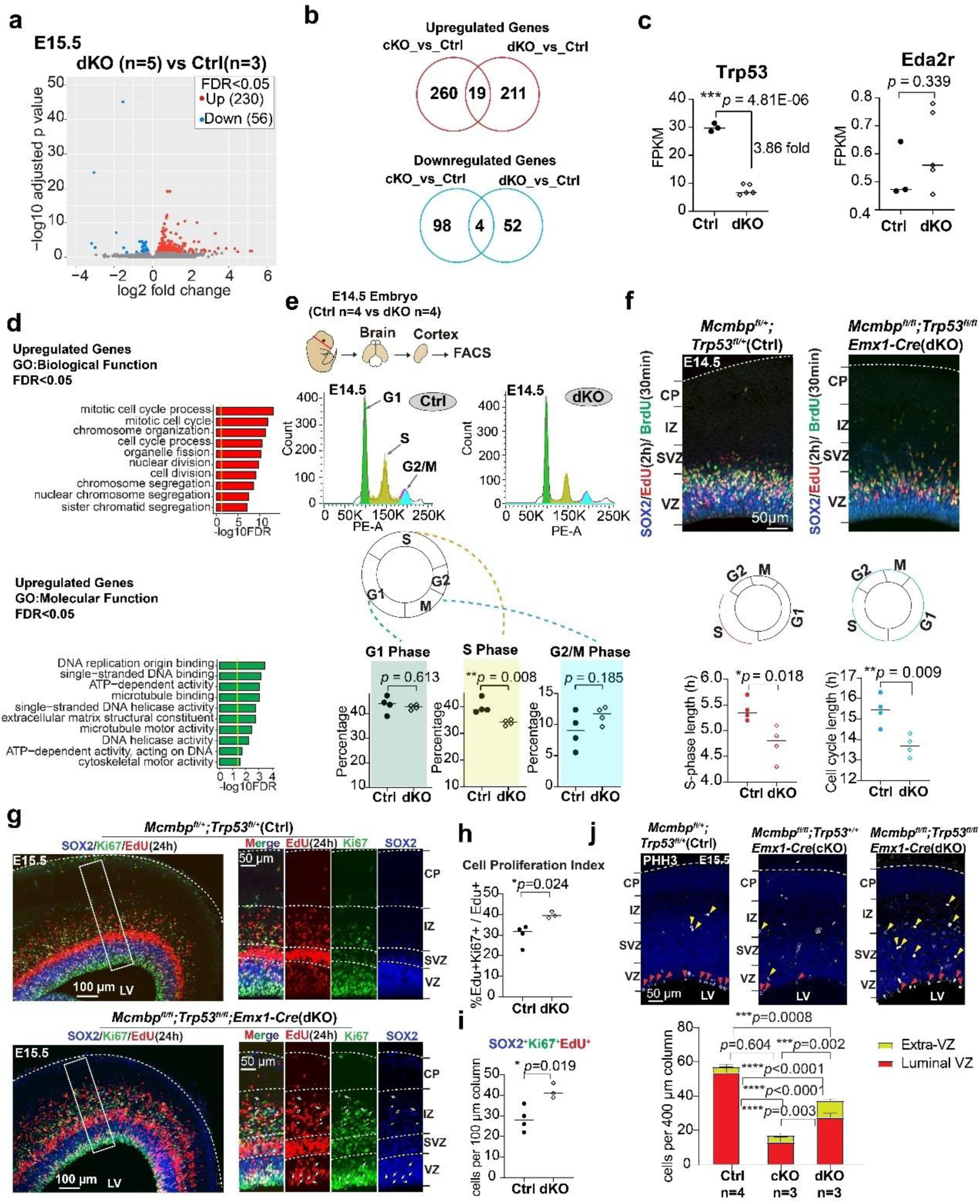
|Cell cycle analysis in *Mcmbp/Trp53* dKO. **a,** Volcano plot showing differentially expressed genes between controls (n=3) and dKO (n=5). Down-regulated genes (FDR<0.05) are indicated in blue and up-regulated genes (FDR<0.05) are indicated in red. **b,** Venn diagram showing small number of overlapping upregulated and downregulated genes between cKO and dKO. **c,** Gene expression plot with FPKM values showing that p53 expression was downregulated in dKO, while its target gene Eda2r expression was similar between dKO (n=5) and controls (n=3). **d,** GO biological and molecular function analysis with upregulated genes. **e,** (Top panel) schematic map of the cell cycle analysis process at E14.5. (Middle panel) cell cycle distribution plots in control and dKO. (Bottom panel) Quantification of proportions of G1, S, and G2/M phase in control and dKO indicating that S phase cells were significantly decreased in dKO. (mean, two-tailed unpaired t-test, ctrl: n=4, dKO: n=4). **f,** Analysis of E14.5 cortex following a 2-hour pulse of EdU (red) and a 0.5-hour pulse of BrdU (green), and quantitative results indicated that both S phase and whole cell cycle length were reduced in dKO. (mean, two-tailed unpaired t-test, ctrl: n=4, dKO: n=4). **g,** Analysis of E15.5 cortex following a 24-hour pulse of EdU (red), arrows indicate SOX2+ NPCs in S phase undergoing proliferation in dKO. **h-i,** Quantification of S phase cells (h) and associated SOX2+ cells(i) between control and dKO. (mean, two-tailed unpaired t-test, ctrl: n=4, dKO: n=3). **j,** Quantification of M phase cells between control, cKO and dKO. (mean, two-tailed unpaired t-test, ctrl: n=4, cKO: n=3, dKO: n=3).

To assess cell cycle dynamics in dKO, we isolated NPCs from the cortex and performed flow cytometry analysis using propidium iodide DNA labeling at E14.5 (**Fig. 5e**). FACS analysis revealed that dKO cells exhibited a selective reduction in S-phase population, while G1 and G2/M phase populations remained comparable to controls (**Fig. 5e**). To precisely quantify cell cycle and S-phase lengths, we performed short-term dual-pulse labeling with EdU (2 hours) and BrdU (30 minutes) at E14.5 (**Fig. 5f**). This analysis confirmed a 12.04% reduction in S-phase length and an 11.21% decrease in total cell cycle duration in dKO (**Fig. 5f**, bottom panel). These findings suggest that accelerated fork speed in dKO may lead to shortened S-phase duration, consequently reducing the proportion of S-phase cells.

Extended analysis using long-term EdU pulse labeling (24 hours) at E15.5 revealed a significantly higher percentage of Ki67+EdU+ proliferating cells compared to controls (**Fig. 5g**, **h**), with SOX2-positive RGCs comprising the majority of this proliferating population (**Fig. 5i**). Notably, we observed detached SOX2-positive RGCs expressing both Ki67 and EdU, indicating maintained proliferative capacity (**Fig. 5i**). Supporting these findings, E15.5 dKO showed increased PHH3-positive cells in extra-ventricular regions (**Fig. 5j**).

Collectively, these results demonstrate that *Mcmbp/Trp53* co-deletion leads to shortened cell cycle duration, particularly in S-phase, potentially enhancing RGC proliferative capacity.

### Single-cell Analysis Reveals Distinct Expression Features of *Trp53/Mcmbp*- deficient RGCs

RGCs are traditionally classified as either apical or basal based on their anatomical location. Basal RGCs, also known as outer radial glial cells (oRGCs), typically reside in the SVZ and are rarely observed in mouse species ^30,31^. This prompted us to investigate whether the displaced progenitors exhibited oRGC-like characteristics. Through double immunostaining for the RGC marker SOX2 and the oRGC marker HOPX ^32^ at E17.5, we identified detached progenitors co-expressing SOX2 and HOPX (**Fig. 6a**).

**Fig. 6.**
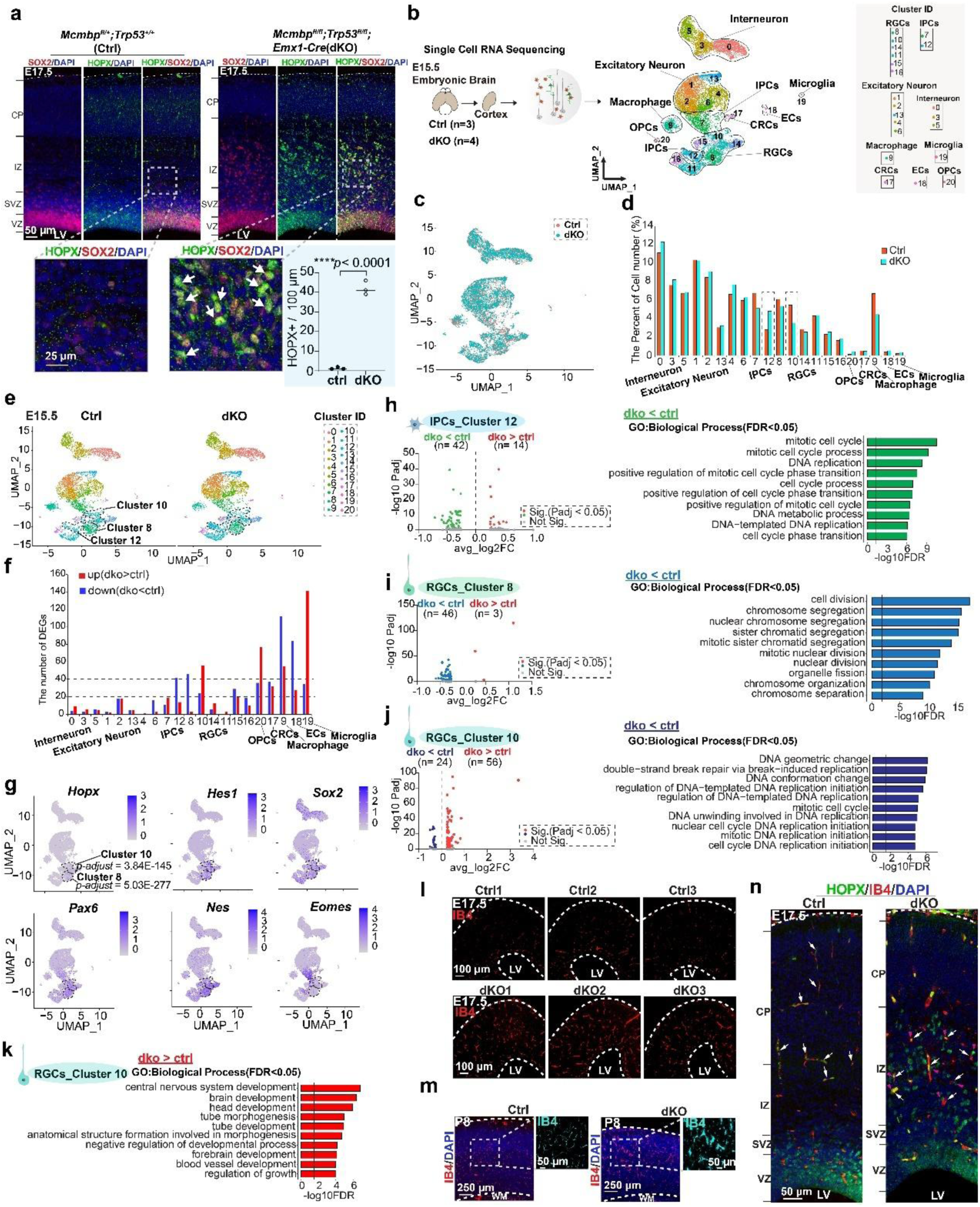
|Single-cell analysis in *Mcmbp/Trp53* dKO identifying expression profile features of progenitors. **a,** Immunostaining of SOX2 and HOPX in E17.5 dKO and their littermate controls. (mean, two-tailed unpaired t-test, ctrl: n=3, dKO: n=3). **b,** Schematic map of single-cell RNA sequencing process and uniform manifold approximation and projection (UMAP) plot showing cell types identified in the integrated controls (n=3) and dKO (n=4) single cell data. **c,** Integrative UMAP between control and dKO. **d,** The percentage of cell number in each identified cluster. **e,** Split UMAP to visualize control and dKO. **f,** The number of upregulated and downregulated genes in each identified cluster. **g,** Feature plots of oRG marker gene *Hopx,* RGC marker genes *Hes1*, *Sox2*, *Nes*, *Pax6*, and IPCs mark gene *Eomes*. **h-j,** (Left panel) volcano plot of DEGs in IPCs_cluster12(h), RGCs_cluster8(i), and RGCs_cluster10(j); (Right panel) GO biological process analysis with downregulated genes in these three clusters. **k,** GO biological process analysis with upregulated genes in RGCs_cluster10. **l-m,** IB4 staining to show blood vessel of cortex in E17.5(l) and postnatal (p8) (m) dKO and their littermate controls. **n,** IB4 and HOPX double staining to show blood vessel associated with HOPX+ progenitors in E17.5 dKO and their littermate controls. White arrows indicate IB4+ cell and pink arrow indicated HOPX+ cells.

To characterize the molecular profile of detached progenitors, we performed single-cell RNA sequencing on control and dKO cortices at E15.5 (**Fig. 6b** and **Extended Data Fig. 7a-c**). Using uniform manifold approximation and projection (UMAP), we identified twenty-one distinct clusters (**Fig. 6b**, left panel) that resolved into nine major cell types based on classical markers: RGCs, IPCs, Cajal-Retzius cells (CRCs), excitatory neurons, interneurons, endothelial cells (ECs), oligodendrocyte progenitor cells (OPCs), macrophages, and microglia (**Fig. 6b**, right panel, and **Supplementary Table 6**). Integrative analysis between control and dKO revealed largely comparable cluster distributions, suggesting successful rescue of *Mcmbp* deletion-induced phenotypes at the single-cell level (**Fig. 6c, d**, and **Supplementary Table 7**). Notably, we observed a 72.67% increase in IPCs_cluster12 population and a 35.71% reduction in RGCs_cluster10 (**Fig. 6d** and **Supplementary Table 7**). Differential expression analysis identified IPCs_cluster12 and RGCs_cluster8 as harboring the highest number of differentially expressed genes within their respective cell types (**Fig. 6e, f**, and **Supplementary Table 7**).

Expression analysis of *Hopx* and other RGC markers revealed significant upregulation of *Hopx* in both RGCs_cluster8 and RGCs_cluster10 (**Fig. 6g**, cluster 8 p-adjust = 5.03E-277; cluster 10 p-adjust = 3.84E-145). Detailed transcriptional analysis of IPCs_cluster12, RGCs_cluster10, and RGCs_cluster8 revealed distinct molecular signatures. IPCs_cluster12 showed 42 downregulated and 14 upregulated genes (**Fig. 6h** and **Supplementary Table 8**), with GO analysis of downregulated genes highlighting enrichment in cell cycle and DNA replication pathways (**Fig. 6h** and **Supplementary Table 8**). In contrast, RGCs_cluster8 exhibited downregulation of genes associated with cell division and nuclear division processes (**Fig. 6i** and **Supplementary Table 9**).

Analysis of the other *Hopx*-positive population, RGCs_cluster10, revealed 80 differentially expressed genes (24 downregulated, 56 upregulated; **Fig. 6j** and **Supplementary Table 10**), exceeding the number found in RGCs_cluster8. Notably, GO analysis of downregulated genes in RGCs_cluster10 showed predominant enrichment in DNA replication-related processes (9/10 biological processes; **Fig. 6j**), strongly suggesting this cluster may represent the detached RGC population. The development pathways (**Fig. 6k** and **Supplementary Table 10**). Using isolectin-IB4 as a blood vessel marker^33^, we confirmed vascular overgrowth in dKO at E17.5 (**Fig. 6l**), with significantly enlarged vessels evident in the dKO cortex by P8 (**Fig. 6m**). Given that blood vessels are essential for RGC proliferation, differentiation, and migration through nutrient and energy provision^34,35^, we performed double staining with IB4 and HOPX. This revealed intimate associations between blood vessels and HOPX-positive progenitors in the dKO cortex, indicating vascular support of these cells (**Fig. 6n**).

### MCM3 Mediates RGC Attachment to the Ventricular Zone

To elucidate the molecular mechanism underlying RGC detachment, we first analyzed MCM2-7 subunit expression through Western blot analysis of whole cell lysates (**Extended Data Fig. 8a**). Consistent with previous studies ^14,18^, MCM2-7 subunit levels were reduced in both cKO and dKO, but not in Emx1-*Trp53* cKO (**Extended Data Fig. 8b**). Given MCMBP’s role in cytoplasmic MCM3-7 assembly, we examined both cytoplasmic and nuclear fractions, finding decreased MCM2-7 subunits in both compartments of cKO and dKO cells, with MCM3 showing unique significant reduction in dKO (**Fig. 7a, b**). Immunostaining of isolated E15.5 NPCs confirmed significantly reduced nuclear MCM3 after *Mcmbp* deletion (**Fig. 7c**).

**Fig. 7.**
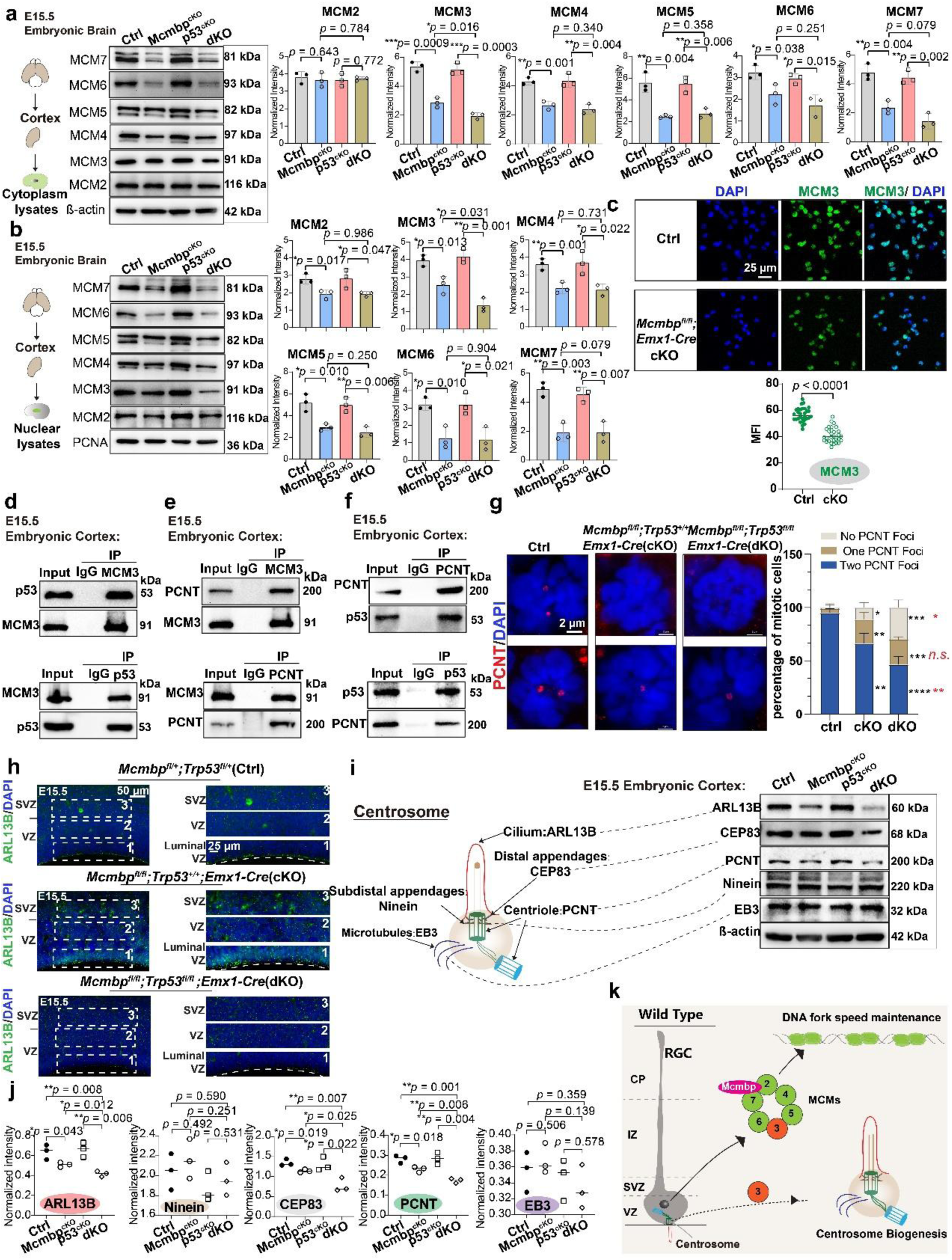
|MCM3 coordinates DNA fork speed and centrosome biogenesis in progenitors. **a,** Western blotting on cytoplasmic lysates and quantitative analysis identifying MCM3 as the only gene significantly repressed in dKO (mean, two-tailed unpaired t-test, ctrl: n=3, *Mcmbp* cKO: n=3, *Trp53*-cKO: n=3, dKO: n=3). **b,** Western blotting on nuclear lysates and quantitative analysis identifying MCM3 as the only gene significantly repressed in dKO. (mean, two-tailed unpaired t-test, ctrl: n=3, *Mcmbp* cKO: n=3, *Trp53*-cKO: n=3, dKO: n=3). **c,** Immunostaining of the MCM3 and quantitative analysis identifying nuclear MCM3 expression was downregulated in cKO. (mean, two-tailed unpaired t-test, ctrl: n=30, cKO: n=30). **d,** Co-immunoprecipitation between MCM3 and p53 in embryonic mouse cortex using MCM3 and p53 as bait, respectively. **e,** Co-immunoprecipitation between MCM3 and PCNT in embryonic mouse cortex using MCM3 and PCNT as bait, respectively. **f,** Co-immunoprecipitation between p53 and PCNT in embryonic mouse cortex using p53 and PCNT as bait, respectively. **g,** (Left panel) centrosome marker PCNT immunostaining in control, cKO and dKO. (Right panel) Quantitative analysis of the percentages of cells with two PCNT foci, one PCNT focus, and no PCNT foci respectively. (mean, two-tailed unpaired t-test, ctrl: 80 cells, cKO: 80 cells, dKO: 80 cells from n=3 samples respectively). **h,** Representative images of the cilium marker ARIL13B at luminal VZ in E15.5 control, cKO, and dKO. **i,** Schematic map of centrosome components and western blotting analysis on their representative markers. **j,** Quantitative analysis of western blotting results for centrosome components. (mean, two- tailed unpaired t-test, ctrl: n=3, *Mcmbp* cKO: n=3, *Trp53*-cKO: n=3, dKO: n=3). **k,** Schematic model of MCM3 coordination of DNA replication and centrosome biogenesis process in radial glial cells.

The specific downregulation of MCM3 in dKO but not cKO suggested potential p53 involvement in MCM3 regulation in RGCs. Co-immunoprecipitation (Co-IP) assays of embryonic cortex extracts confirmed *in vivo* interaction between p53 and MCM3 (**Fig. 7d**). While MCM subunits are known to participate in centriole duplication, which mediates RGC anchoring to the ventricular surface^36–38^, MCM3’s specific role in RGC centrosome biogenesis remained unclear. Co-IP assays revealed interactions between MCM3 and the centrosome marker pericentrin (PCNT) (**Fig. 7e**), and between PCNT and p53 (**Fig. 7f**). These findings suggest that MCM3 coordinates DNA and centriole replication during neurogenesis through protein-protein interactions.

In wild-type cells, the majority of mitotic cells at the VZ surface contained two PCNT foci (95%, compared to 3.75% with one focus and 1.25% with none; **Fig. 7g**). However, in cKO, only 66.25% of mitotic cells maintained two PCNT foci, with significant increases in cells containing one (22.50%) or no (11.25%) foci. This phenotype was further exacerbated in dKO, where only 23.75% of cells retained two PCNT foci, with dramatic increases in cells showing one (23.75%) or no (30%) foci (**Fig. 7g**), indicating that *Trp53* deletion aggravated centrosome duplication defects following *Mcmbp* deletion.

ARL13B, a primary cilia marker and essential component for polarized radial glial scaffold formation ^39^, showed uniform distribution along the VZ surface in controls. In contrast, cKO exhibited disrupted distribution patterns, with further dramatic reduction in dKO (**Fig. 7h, i**), indicating compromised centrosome integrity. Given that centrosomes serve as major microtubule-organizing centers (MTOCs) essential for cell division and comprise multiple components (cilium, distal appendages, subdistal appendages, centriole, and microtubules; **Fig. 7i**), we analyzed specific centrosomal components by Western blot (**Fig. 7i**). Quantification revealed downregulation of multiple components in both cKO and dKO: CEP83 (distal appendage), PCNT (centriole), and ARL13B (cilium), with p53 deletion enhancing these effects (**Fig. 7j**). These findings indicate progressive RGC detachment from the VZ surface due to selective disruption of centrosome components.

The VZ provides a specialized niche for RGCs, where apical-basal polarity and surface anchoring depend on both centrosome integrity and adhesion molecules such as N-cadherin (NCad). NCad, a calcium-dependent adhesion molecule, mediates both homotypic and heterotypic cell-cell interactions ^40^. Notably, dKO showed reduced NCad immunostaining at the VZ surface (**Extended Data Fig. 8c**), suggesting that compromised cell-cell interactions also contribute to RGC detachment.

In summary, wild-type RGCs utilize MCMBP to facilitate nuclear import of newly formed MCM2-7 complexes, which regulate DNA fork speed as pre-RC components, while a subset of MCM3 participates in centrosome biogenesis (**Fig. 7k**).

### Disruption of Fork Speed Induces Anxiety-like Behavior in Adult Mice

To investigate the long-term consequences of disrupted DNA fork speed in RGCs, we examined adult mouse behavior. At postnatal day 45 (P45), dKO brain morphology appeared comparable to controls (**Extended Data Fig. 9a**). Analysis of the corpus callosum (CC), the largest white matter fiber tract connecting the hemispheres^41^, revealed thinning and bifurcation in cKO brains, while dKO showed rescued CC morphology similar to controls (**Extended Data Fig. 9b**).

Behavioral analysis of three-month-old mice using a 3D Mouse behavior system ^42^ revealed distinct phenotypes. Principal component analysis (PCA) of kinematic features showed clear separation of the cKO group from control and dKO groups (**Fig. 8a**). The dKO group clustered near but slightly separate from controls, suggesting partial behavioral restoration accompanying the rescued brain size (**Fig. 8a**). Open field trajectories further confirmed the similarity between dKO and control patterns compared to cKO (**Fig. 8b**). However, both cKO and dKO mice exhibited reduced time spent in the open field center relative to controls (**Fig. 8c**), indicating persistent anxiety-like behavior despite the rescue of brain size, CC integrity, and other structural features. Detailed phenotypic analysis using the 3D Mouse system revealed multiple behavioral dimensions. Regarding body morphology, dKO mice showed control-like patterns in body length and angle measurements, both exceeding cKO values (**Extended Data Fig. 9c**). Natural behaviors showed striking differences: cKO mice exhibited increased grooming frequency with reduced sniffing, rising, and rearing behaviors compared to controls (**Extended Data Fig. 9d**). While dKO mice showed normalized grooming and rising behaviors, they maintained altered sniffing (decreased) and rearing (increased) frequencies (**Extended Data Fig. 9d**), indicating persistent behavioral abnormalities. Movement parameters, including tail speed and tail tip energy, showed strong similarity between dKO and controls (**Extended Data Fig. 9e**).

**Fig. 8.**
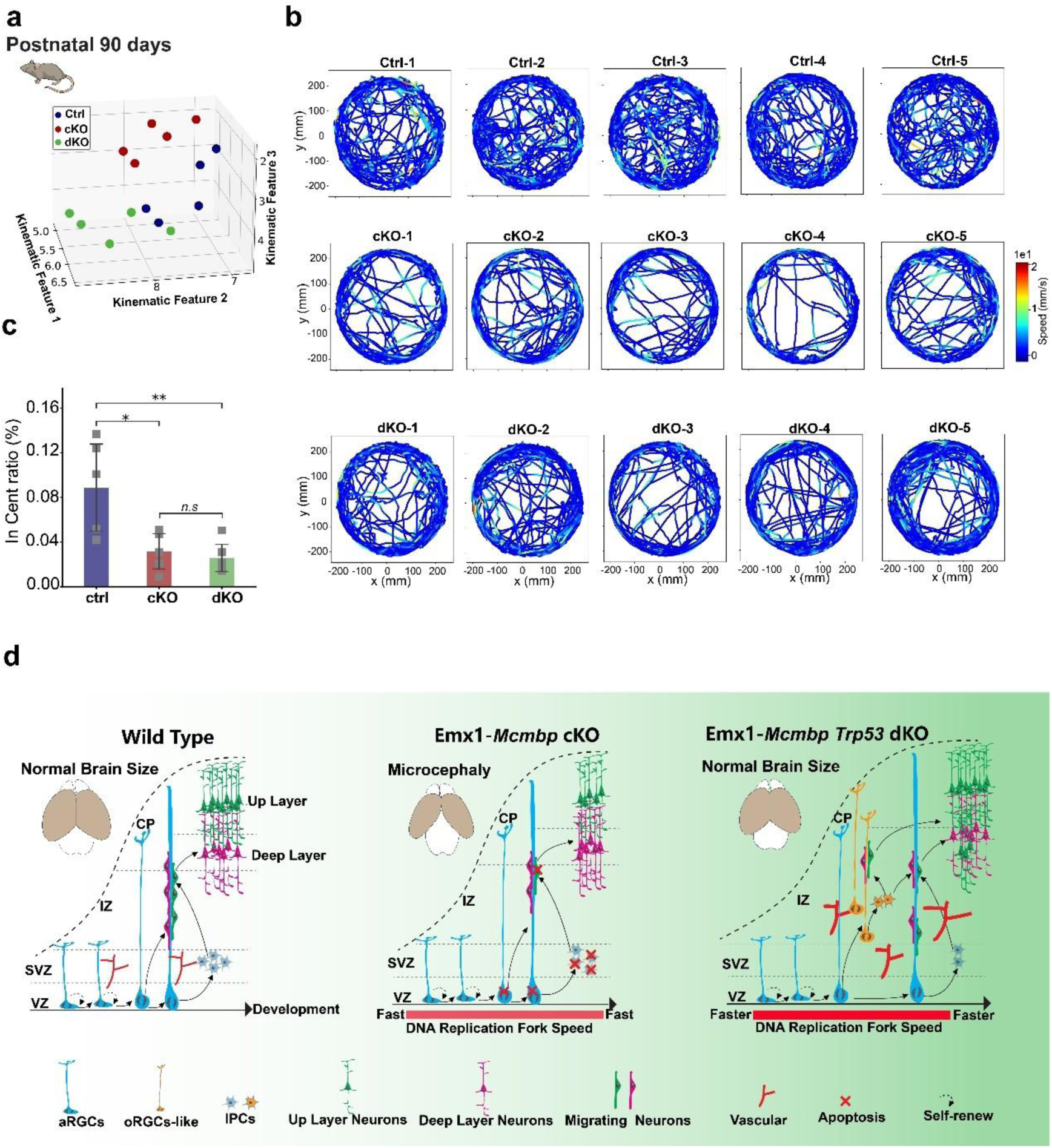
|Fork speeding acceleration leads to an anxiety-like behavior in *Mcmbp* cKO and *Mcmbp/Trp53* dKO. **a,** PCA analysis of kinematic features in controls, cKO, and dKO. **b,** Track records of open field assay from 5 controls, 5 cKO, and 5 dKO. **c,** Bar plots of center-to-total area ratio in controls, cKO, and dKO. (Welch’s t-test, ctrl, cKO, dKO: n=5). **d,** The proposed model for DNA fork speed regulation during neurogenesis. During cortical development, DNA fork speed progression is restrained through the DNA replication complex MCMs, which are assembled, stabilized and exported by MCMBP. In contrast to fork speed progression, the expression of MCMBP is gradually decreased throughout the neurogenesis. Consequently, after deletion of *Mcmbp* in the early stages using Emx1-Cre, fork speed was increased (light red) in RGCs, which leads to massive cell apoptosis, and results in a microcephaly phenotype. In the absence of *Mcmbp* and *Trp53*, although the brain size is largely rescued, DNA fork speed was unexpectedly increased (red color) and faster than deletion of *Mcmbp* alone, resulting in many RGCs detached from the VZ and became oRG-like cells (orange color). Meanwhile, the blood vessel was overgrown in dKO (*Mcmbp/Trp53* co- deletion).

Collectively, these findings demonstrate that disruption of fork speed during cortical development impacts brain maturation and produces lasting behavioral alterations in adulthood, even when gross anatomical features are restored.

## Discussion

In cortical neurogenesis, genome stability and integrity are fundamental prerequisites for proper brain development ^43^. DNA replication serves as the cornerstone of genome duplication in neural stem cells during their proliferation and differentiation into daughter cells. Consequently, perturbations in genes involved in DNA duplication processes during neurogenesis can result in neurodevelopmental disorders. For instance, the *Topbp1* gene, essential for DNA replication firing, leads to severe defects in mammalian cortical and cerebellar development when absent ^44^. Similarly, *DONSON*, identified as a critical component of CMG helicase assembly ^45^, causes microcephalic dwarfism in humans when mutated ^46,47^. However, the specific role of DNA replication fork speed in regulating radial glial cell (RGC) fate and its implications for brain development remained unexplored.

Our study identifies DNA fork speed as a crucial regulator of RGC fate. We observed that MCMBP exhibits high expression during early developmental stages, followed by a significant decline during later division phases (**Fig. 1b, c**). To investigate the role of DNA fork speed in neurogenesis, we generated *Mcmbp* cKO mice. Our findings revealed that *Mcmbp* deletion resulted in increased DNA fork speed in RGCs, triggering widespread DNA damage, p53 activation, and extensive cell apoptosis, ultimately manifesting as a microcephaly phenotype. Intriguingly, while concurrent deletion of both *Mcmbp* and *Trp53* largely rescued brain size, it led to even faster DNA fork speeds than *Mcmbp* deletion alone, causing RGCs to detach from the ventricular zone and acquire outer RGC-like characteristics. These results strongly support a model where DNA fork speed functions as a cellular pacer that requires precise regulation during cortical neurogenesis (**Fig. 8d**). Furthermore, recent studies have demonstrated that reducing DNA fork speed can modulate cellular plasticity and enhance the reprogramming efficiency of pluripotent stem cells into totipotent-like cells^48^. This raises intriguing questions about the potential effects and underlying mechanisms of decreased fork speed in RGCs during cortical development.

Mechanistically, MCMBP functions as a critical chaperone protein that facilitates the assembly, stabilization, and translocation of nascent MCMs, thereby regulating DNA fork speed to maintain genome integrity ^14^. In RGCs lacking *Mcmbp*, MCM subunit expression is downregulated, leading to dysregulated DNA fork speed. Notably, simultaneous deletion of both *Mcmbp* and *Trp53* results in even greater acceleration of fork speed compared to *Mcmbp* deletion alone. Our co-immunoprecipitation studies revealed a direct interaction between p53 and MCM3, suggesting that *p53* may protect MCM3 from degradation in MCMBP’s absence. This finding presents an interesting contrast to previous research indicating that *Trp53* deletion typically reduces DNA fork progression ^49,50^.

The coordination between centrosome duplication and DNA replication is essential for cell cycle progression ^51^, though the precise mechanisms governing this synchronization remain elusive. Our observations demonstrate that *Mcmbp* deletion disrupts both centrosome number and components, indicating an intricate relationship between fork speed regulation and centrosome biogenesis during the cell cycle. Significantly, we found that both *p53* and MCM3 interact with the centrosomal protein PCNT, suggesting a molecular link between DNA replication completion in S phase and subsequent centrosome duplication for chromosome segregation. This relationship extends to RGC division orientation, as centrosomes play a crucial role in determining mitotic spindle direction ^52,53^.

An unexpected finding was that RGCs exhibiting accelerated fork speed persisted in dKO mice but dispersed into extra-ventricular zone regions during brain development. This phenotype parallels observations in *Sas-4* (*Cenpj*)/*Trp53* double knockouts, where RGCs similarly delocalize to extra-VZ regions ^38^. However, a key distinction emerges: while *Sas-4*-induced ectopic RGCs maintain limited proliferative and differentiative capabilities similar to apical RGCs, the displaced RGCs resulting from *Mcmbp/p53* deletion exhibit enhanced proliferation and strong expression of the oRGC marker HOPX. These characteristics mirror those of human outer radial glial cells, particularly their elevated proliferative capacity ^54^—a trait considered crucial in the evolutionary expansion of human brain size relative to chimpanzees ^55^.

The observed blood vessel overgrowth suggests that maintaining these highly proliferative RGCs requires sustained energy input. Intriguingly, while angiogenesis typically proceeds independently and only responds to RGC-derived VEGF signaling in later stages ^56^, vascular structures show preferential contact with proliferative cells to modulate cell cycle processes ^34^. This raises important questions about the mechanisms by which vascular filopodia interact with displaced RGCs exhibiting accelerated fork speed, warranting further investigation.

In conclusion, while our study did not explore the functional effects of other fork speed-regulating genes in neurogenesis, we have uncovered a novel mechanism whereby DNA fork speed orchestrates RGC fate and behavior through coordinated regulation of DNA replication and centrosome biogenesis. This discovery significantly advances our understanding of DNA fork speed’s role in cellular plasticity regulation.

## Methods

### Ethics Statement

All mouse experiments were conducted in accordance with guidelines approved by the Animal Care and Use Committee of the Kunming Institute of Zoology, Chinese Academy of Sciences. Animals were maintained in standard feeding environments and were free of pathogens or parasites that could significantly impact experimental outcomes. *Mcmbp^fl/fl^* mice were obtained from Jiangsu Gempharmatech Co., Ltd. C57BL/6J mice and *Emx1-Cre*^21^ were sourced from Jackson Laboratory (Bar Harbor, ME, USA). Wild-type C57BL/6J mice were acquired from the Animal Center of Kunming Institute of Zoology, Chinese Academy of Sciences.

### Generation of *Mcmbp* cKO mice

The *Mcmbp* gene comprises five transcripts. Based on the gene structure analysis, we targeted exons 6-8 of the *Mcmbp*-201 transcript (ENSMUST00000057557.13) for knockout, encompassing a 398bp coding sequence. Disruption of this region using CRISPR/Cas9 technology results in loss of protein function. The generation process, illustrated in **Extended Data Fig. 1b**, involved *in vitro* transcription of sgRNA and construction of the donor vector. The Cas9 protein, sgRNA, and donor constructs were microinjected into C57BL/6J fertilized eggs, which were subsequently transplanted into recipient females. Positive F0 mice were identified through PCR and sequencing, then bred with C57BL/6J mice to generate F1 offspring. *Mcmbp* cKO mice were produced by crossing *Mcmbp^fl/fl^* mice with *Emx1-Cre* mice and confirmed through genotyping. Oligonucleotide primers used for genotyping are detailed in **Supplementary Table 11**.

### Isolation and Culture of Mouse Neural Progenitor Cells

Neural progenitor cells were isolated as single-cell suspensions from mouse cerebral cortex and maintained in proliferation medium comprising NeuroCult™ Basal Medium (Stemcell, 05700), NeuroCult™ Proliferation Supplement (Stemcell, 05701), 2% B27 supplement (Gibco, 12587010), 20 ng/ml epidermal growth factor (EGF, STEMCELL, 78006), 10 ng/ml basic fibroblast growth factor (bFGF, STEMCELL, 78003), 1% penicillin/streptomycin (100 units/mL penicillin and 100 µg/mL streptomycin) (Gibco, 15140-122), and 0.0002% heparin (STEMCELL, 07980). Cells were enzymatically dissociated using papain (Worthington, Ls00319) during passaging and cultured on 6- well plates pre-coated with 5µg/mL laminin (Invitrogen, 23017015) and 10 µg/mL poly-d-lysine (Sigma, P3655).

### Immunofluorescence Staining and Imaging

Brain tissues were harvested at various developmental stages and fixed in 4% paraformaldehyde (Acros Organics, AC416785000) for 24 hours at room temperature. Tissues were embedded in 4% agarose and sectioned at 35-70 μm thickness using a Vibratome (Leica, VT1000S). For immunostaining, antigen retrieval was performed using citrate buffer (Servicebio, G1202). Sections underwent three PBS washes followed by permeabilization in PBS containing 0.5% Triton X-100 (MACKLIN, TB24275) for 30 minutes. Sections were then blocked for 1-2 hours at room temperature in PBS containing 5% normal donkey serum (Jackson, 017-000-121), 1% bovine serum albumin (Sigma, SRE0096), 0.1% glycine (BBI, A610235-0500), and 0.1% lysine (MACKLIN, L812314). Primary antibody incubation was performed overnight at 4°C in blocking buffer. Following three 5-minute PBS washes, sections were incubated with fluorescent secondary antibodies for 2 hours at room temperature, protected from light. Nuclei were counterstained with DAPI (1:1000, Invitrogen, 62248). Sections were mounted using VECTASHIELD Antifade Mountant (Vector Labs, H-1000). Images were acquired on a Leica TCS SP8 confocal microscope and analyzed using ImageJ software (National Institutes of Health, NIH).

### Primary Antibodies

Mouse Monoclonal anti-BRN2/POU3F2 (1:250, Santa Cruz Biotechnology, sc- 393324); Rabbit Polyclonal anti-TBR1 (1:250, Affinity, DF2396); Rabbit Polyclonal anti-EOMES (1:500, Millipore, ABE1402); Mouse Monoclonal anti-TLE4 (1:100, Santa Cruz Biotechnology, sc-365406); Rat Monoclonal anti-SOX2 (1:500, Invitrogen, 53-9811-82); Rabbit Polyclonal anti-TBR2 / EOMES (1:500, Abcam, ab23345); Phospho-Histone H3 (Ser10) Antibody (1:500, Cell Signaling Technology, 9701S); Rabbit Monoclonal anti-Ki67 (1:500, Cell Signaling Technology, 9129S); Rabbit Monoclonal anti-P53(7F5) (1:500, Cell Signaling Technology, 25247S); Rabbit Monoclonal anti- Phospho-Histone H2A.X(Ser139) (1:500, Cell Signaling Technology, 9718S); Rabbit Polyclonal anti-Cleaved Caspase-3 (Asp175) (1:500, Cell Signaling Technology, 9661S); Rabbit Polyclonal anti-Phospho-KAP-1(Ser824) (1:500, Bethyl, A300-767A); Mouse Monoclonal anti-N-cadherin (1:250, BD, 610920); Mouse Monoclonal anti-BrdU (1:250, Invitrogen, B35128); Rabbit Polyclonal anti-MCM3 (1:250, Santa Cruz Biotechnology, SC-390480); Mouse Monoclonal anti-p- vimentin(1:500, Enzo, ADI-KAM-CC249-E); Rabbit Polyclonal anti-ARL13B (1:250, Proteintech, 66009-1-Ig); Rabbit Polyclonal anti-Pericentrin(1:250, Abcam, ab4448); Rat Monoclonal anti-Neural Cell Adhesion Molecule L1(L1CAM) (1:500, Millipore, MAB5272); Secondary antibodies: Alexa Fluor 488 AffiniPure Donkey Anti-Rabbit (1:500, Jackson, 711-545-152); Alexa Fluor 488 AffiniPure Donkey Anti-Rat (1:500, Jackson, 712-545-150); Alexa Fluor 488 AffiniPure Donkey Anti-Mouse (1:500, Jackson, 715-545-150); Alexa Fluor 594 AffiniPure Donkey Anti-Rabbit (1:500, Jackson, 711-585-152); Alexa Fluor 594 AffiniPure Donkey Anti-Rat (1:500, Jackson, 712-585-153); Alexa Fluor 594 AffiniPure Donkey Anti-Mouse (1:500, Jackson, 715-585-150).

### Protein Extraction from Cytoplasmic and Nuclear Fractions

Cytoplasmic and nuclear proteins were extracted from E15.5 mouse embryonic cortex using the Beyotime kit (P0027). Fresh cortical tissue was processed with a 20:1 mixture of Cytoplasmic Protein Extraction Reagents A and B. Following 15-minute incubation on ice, the mixture was centrifuged at 15,000 x g for 5 minutes at 4°C to obtain the initial cytoplasmic protein fraction. The pellet was resuspended in 200 µl of Cytoplasmic Protein Extraction Reagent A containing PMSF and incubated on ice for 10-15 minutes. After adding 10 µl of Cytoplasmic Protein Extraction Reagent B, the sample was incubated on ice for 1 minute and centrifuged at 15,000 x g for 5 minutes at 4°C. The supernatants from both centrifugation steps were pooled to obtain the total cytoplasmic protein fraction.

For nuclear protein extraction, the remaining pellet was thoroughly cleared of residual supernatant and resuspended in 50 µl of Nuclear Protein Extraction Reagent containing PMSF. The suspension was vortexed at maximum speed for 15-30 seconds to ensure complete pellet dispersion, followed by a 30-minute incubation on ice with 15-30 seconds of vortexing every 3 minutes. The sample was then centrifuged at 15,000 x g for 10 minutes at 4°C, with the resulting supernatant containing the nuclear protein fraction.

For total protein extraction, fresh cortical tissue was homogenized in RIPA buffer (Beyotime, P0013B) supplemented with protease inhibitor cocktail (Beyotime, P1005) and phosphatase inhibitor (Beyotime, P1082). The tissue was sonicated at 4°C for 5 cycles of 10 seconds each, followed by incubation on a shaker at 4°C for 10 minutes. After centrifugation at 15,000 x g for 10 minutes at 4°C, the supernatant was collected as the total protein solution.

### Western Blotting

Protein samples were resolved by 10% sodium dodecyl sulfate-polyacrylamide gel electrophoresis (SDS-PAGE) and transferred to PVDF membranes (Millipore, IPVH00010). Membranes were blocked with 5% skimmed milk (BBI, A600669-0250) and incubated with primary antibodies overnight at 4°C. Following TBST washes, membranes were incubated for 1 hour with horseradish peroxidase-conjugated secondary antibodies: anti-mouse (1:1000, Proteintech, SA00001-1) and anti-rabbit (1:1000, Proteintech, SA00001-2). Protein bands were visualized using the SuperSignal West Pico PLUS chemiluminescence kit (Thermo Fisher Scientific, 34579) according to manufacturer’s specifications.

The following primary antibodies were used for immunoblotting: MCMBP (1:2000, Invitrogen, PA5-58410), MCM2 (1:1000, Proteintech, 66009-1-Ig), MCM3 (1:1000, Santa Cruz Biotechnology, SC-390480), MCM4 (1:1000, Proteintech, 13043-1-AP), MCM5 (1:1000, Proteintech, 11703-1-AP), MCM6 (1:1000, Santa Cruze Biotechnology, SC-393618), MCM7 (1:1000, Proteintech, 11225-1-AP), ARL13B (1:1000, Proteintech, 17711-1-AP), CEP83(1:1000, Proteintech, 26013-1-AP), PCNT (1:2000, Abcam, ab220784), NINEIN (1:1000, Santa Cruze Biotechnology, SC376420), EB3 (1:1000, 1:1000, Proteintech, 23974-1-AP), β-actin (1:2000, Proteintech, 66009-1-Ig), PCNA (1:1000, Servicebio, GB12010).

### Co-immunoprecipitation Assays

Co-immunoprecipitation (Co-IP) was performed using fresh cortical tissue lysed in buffer containing proteinase inhibitor cocktail (Beyotime, P1005). Following 30- minute incubation on ice, the lysate was centrifuged at 12,000 rpm for 10 minutes at 4°C, and the supernatant was collected. Fifty microliters of Protein A/G magnetic beads (Medchemexpress, HY-K02002) were used for magnetic separation, and the supernatant was removed. The beads were resuspended in solution containing primary antibodies against PCNT (Abcam, ab220784), MCM3 (Santa Cruz, SC-390480), and P53 (CST, 25247S) at 20 µg/ml concentration and incubated with rotation overnight at 4°C.

Following magnetic separation and supernatant removal, the beads were resuspended in 50 µg of protein lysate and incubated with rotation overnight at 4°C. After a final magnetic separation and supernatant removal, the beads were resuspended in SDS loading buffer. Proteins were denatured by heating at 95°C for 7 minutes and analyzed by 10% SDS-PAGE.

### Real-time Quantitative RT-PCR

Total RNA was isolated from mouse cerebral cortex using the TIANGEN extraction kit (DP419). First-strand cDNA synthesis was performed using the HiScript III All-in-one RT SuperMix Kit (Vazyme, R333-01). Quantitative real-time PCR was conducted using the AceQ Universal SYBR qPCR Master Mix Kit (Vazyme, Q511-02) on an Applied Biosystems 7500 Real-Time PCR System (Waltham, MA, USA). Gene expression was normalized to 18sRNA as the reference gene, with normalized values expressed as relative quantity of *Eda2r* mRNA. Oligonucleotide primer sequences used for qRT-PCR amplification are provided in **Supplemental Table 12**.

### Cell Cycle Analysis

Neural progenitor cells were isolated from E14.5 embryonic mouse cortex and fixed in pre-cooled 70% ethanol, followed by 48-hour incubation at 4°C on a rocking platform. After centrifugation at 1000 rpm to remove ethanol, cells were washed twice with pre-cooled PBS. Cells were then incubated in the dark at 4°C for 30 minutes with a solution containing 50 μg/mL propidium iodide (Sigma, P4170), 0.2% Triton X-100, and 100 μg/mL RNase A (Thermo Fisher Scientific, EN0531). Analysis was performed using a Becton-Dickinson LSRFortessa flow cytometer (Franklin Lakes, NJ, USA), and data were analyzed using FlowJo software (FlowJo, LLC, Ashland, OR, USA).

### Cell Cycle Kinetics Calculation

Cell cycle kinetics were assessed using the bis-thymidine labeling assay. At E14.5 (T=0 h), pregnant mice were injected with EdU (10 mg/ml, 100 μl/100 g body weight) to label all S-phase cells. At T=1.5 h, mice received BrdU injection (1 mg/ml, 100 μl/100 g body weight) to label cells in S-phase at the experiment’s conclusion. Animals were euthanized at T=2 h and embryos were collected immediately ^57^. Embryo sections were immunostained using anti-BrdU antibody (Invitrogen, ab6326) and EdU staining solution (100 μL 1M Tris, 100 μL 1M ascorbic acid, 794 μL PBS, 4 μL 1M CuSO4, and 2 μL 4 mM Sulfo-Cyanine3 azide; Lumiprobe, A1330). S-phase duration (Ts) and total cell cycle length (Tc) were calculated using the following equations: *T*_s_ = 1.5 h/(*N*_Edu+Brdu−_/*N*_EdU+BrdU+_) and *T*_c_ = *T*_s_/(*N*_EdU+BrdU+_/*N*_total_) where N represents cell numbers in the respective populations.

### DNA Fiber Assay

Neural progenitor cells were sequentially labeled with 50 μM IdU and 50 μM CldU for 30 minutes each. After centrifugation, cells were resuspended in pre-cooled PBS at 1 ×10^5 cells/ml. Two microliters of cell suspension were placed at one end of a microscope slide, followed by addition of 10 μL lysis buffer (0.5% SDS, 50 mM EDTA, 200 mM Tris-HCl, pH 7.4). The slide was tilted at 15°to ensure uniform flow of the droplet. Cells were fixed with methanol-acetic acid (3:1) and DNA was denatured in 2.5 M HCl at 37°C for 20 minutes.

Following 1-hour incubation with 1% BSA at room temperature, slides were incubated overnight at 4°C with primary antibodies: Mouse Monoclonal anti-BrdU (1:100, BD Biosciences, 347580) and Rat Monoclonal anti-BrdU (1:100, Abcam, ab6326). Secondary antibody incubation was performed for 2 hours at room temperature using Alexa Fluor 488 AffiniPure Donkey Anti-Mouse (1:500, Jackson, 715-545-150) and Alexa Fluor 594 AffiniPure Donkey Anti-Rat (1:500, Jackson, 712-585-153).

Images were acquired using a confocal microscope and analyzed with ImageJ software. DNA fiber length measurements were calibrated using the conversion factor of 2.59 kb per 1 μm of spread DNA ^58^. Statistical analysis was performed using Student’s t-test for two-group comparisons or one-way ANOVA for multiple group comparisons.

### RNA Sequencing Library Construction and Illumina Sequencing

RNA purification, reverse transcription, library construction, and sequencing were performed at Sequanta Technologies Co., Ltd (Shanghai) following manufacturer’s protocols. Globin mRNA was depleted using GLOBINclear™ Mouse/Rat Kit (Invitrogen, AM1981). Non-Globin mRNA sequencing libraries were prepared from 1 µg total RNA using VAHTS mRNA-seq v3 Library Prep Kit (Vazyme, NR611).

PolyA mRNA was isolated from total RNA using oligo-dT-attached magnetic beads and fragmented using fragmentation buffer. These fragments served as templates for first-strand cDNA synthesis using reverse transcriptase and random primers, followed by second-strand synthesis. The resulting cDNA underwent end-repair, phosphorylation, and ’A’ base addition according to Illumina’s library construction protocol. Illumina sequencing adapters were then ligated to both ends of the cDNA fragments.

Following PCR amplification for DNA enrichment, target fragments of 200-300 bp were purified using AMPure XP Beads (Beckmen, A63880). Library quality control was performed using Qubit 3.0 fluorometer dsDNA HS Assay (Thermo Fisher Scientific) for concentration measurement and Agilent BioAnalyzer (Being, CHN) for size distribution analysis. Sequencing was conducted on an Illumina Novaseq 6000 platform using 2 x 150 paired-end sequencing protocols at Sequanta Technologies Co., Ltd.

### Bulk RNA-seq Data Analysis

(https://www.bioinformatics.babraham.ac.uk/projects/fastqc/), followed by adapter sequence trimming using Cutadapt tools^59^. The processed reads were mapped to the mouse reference genome (mm10) using HISAT2 2.1.0^60^. Gene expression quantification was performed using HTSeq-count (http://www-huber.embl.de/users/anders/HTSeq/doc/count.html) to obtain integer counts of mapped reads per gene.

FPKM expression values were calculated using Cufflinks v2.2.1^61^, incorporating automatic estimation of library size distributions and sequence composition bias correction. Differential expression analysis was conducted using R package DESeq2 version 1.36.0^62^, which employs negative binomial distribution modeling of count data. Volcano plots were generated using the DESeqAnalysis R package^63^.

Gene ontology (GO) enrichment analysis of differentially expressed genes was performed using the ToppGene Suite^64^. Only functional annotation terms showing significant enrichment (Bonferroni correction, p<0.05) compared to the total gene population were included in the clustering analysis. GO terms were analyzed for both biological processes and molecular functions.

### Single-cell RNA Capture, Reverse Transcription and Library Construction

Single-cell RNA processing was performed using SeekOne® Digital Droplet (SeekOne® DD) system and associated kits from SeekOne Bio, including: SeekOne® DD S3 Microarray Kit (SEEKGENE, K00202-0201), SeekOne® DD Single-Cell Purification Kit (SEEKGENE, K00202-0205), SeekOne® DD Single-Cell Whole-Sequence Microbead Kit (SEEKGENE, K00801-0202), SeekOne® DD Single-Cell Full Sequence Reverse Transcription Kit (SEEKGENE, K00801-0203) and SeekOne® DD Library Construction Kit (SEEKGENE, K00202-0204). The SeekOne® DD platform enables comprehensive processing from single-cell nucleic acid labeling to full-sequence transcriptome library construction. The system employs microfluidic-based water-in-oil droplet technology for single-cell isolation and capture. RNA molecules from individual cells are labeled using nucleic acid-modified Barcoded Beads, and full-length coding and non-coding RNAs are captured using random primers. Intracellular rRNA and mitochondrial contamination are eliminated using proprietary blocker technology. Libraries were constructed following the “SeekOne® DD Single-Cell Whole Sequence Transcriptome Kit Instruction Manual” to ensure compatibility with high-throughput sequencers. Quality control criteria for qualified libraries included: main peak fragment size range: 450-900 bp, absence of small fragments (verified by Agilent 4200 TapeStation), Minimum library concentration: 1 ng/μL (measured by Qubit 4.0). If small fragments were detected, additional 0.6x purification steps were performed until complete removal was achieved.

### Single-cell Data Analysis

Single-cell data analysis was performed using the Seurat R package^65^. Feature barcode matrices were processed and analyzed using the Seurat RunUMAP function for nonlinear dimension reduction. Comparative analysis between control and *Mcmbp/Trp53* dKO samples was conducted using the FindIntegrationAnchors function, which identifies anchors representing cells with similar biological states based on canonical correlation analysis^66^. Integrated analyses were then performed on all cells following the standard Seurat package workflow.

### Behavioural Experiments

Locomotor activity and anxiety-like behavior were assessed in 3-month-old male mice using the Open Field Test ^67^. Testing was conducted using the BAYONE BIOTECH BehaviorAtlas Analyzer-3D-Mouse Behavioural Device, which enables multi-viewpoint video acquisition ^42^. The testing arena consisted of a circular open field with a white plastic floor and transparent acrylic walls (base diameter: 50 cm; wall height: 30 cm). Mice were given 7 minutes for free exploration, followed by 20 minutes of locomotor activity recording. The arena was cleaned between trials. Behavioral data were recorded using BehaviorAtlas Explorer visualization software and analyzed using BehaviorAtlas Analyzer-Mouse analysis software.

## Data Availability

All sequencing data were deposited at the NCBI Gene Expression Omnibus (GEO) under accession number GSE279962

## Quantification and Statistical Analysis

The *t-test* was used to determine the significant difference by comparing means between two groups of data. Paired data were from the same group but under different conditions was performed using an unpaired or paired two-tailed *Student’s t-test*.

## Code availability

No custom code was generated for this study. All analyses were performed using commercially available software packages, and relevant code is available upon request.

## Acknowledgements

We thank the members of the Shi Laboratory for their discussions and comments on the manuscript. We would like to thank the Core Technology Facility of the Kunming Institute of Zoology, CAS for providing flow cytometry analysis and sorting. We are grateful to Guolan Ma and Shuangjuan Yang for providing technical support. We also thank Shenzhen Bayone BioTech Co., Ltd for providing the BehaviorAtlas Analyzer- 3D Mouse Behavioral Device and technical support.

## Author contributions

L.S. conceived and designed the study; JH.W. performed mouse experiments, JH.W., YF. K., XZ. L., YL. T, and K.X. performed molecular biology experiments; L.S. performed bioinformatics analysis; L.S., and JH.W. prepared the figures; L.S., and JH.W. wrote the final manuscript.

---

**Extended Data Fig.1.**
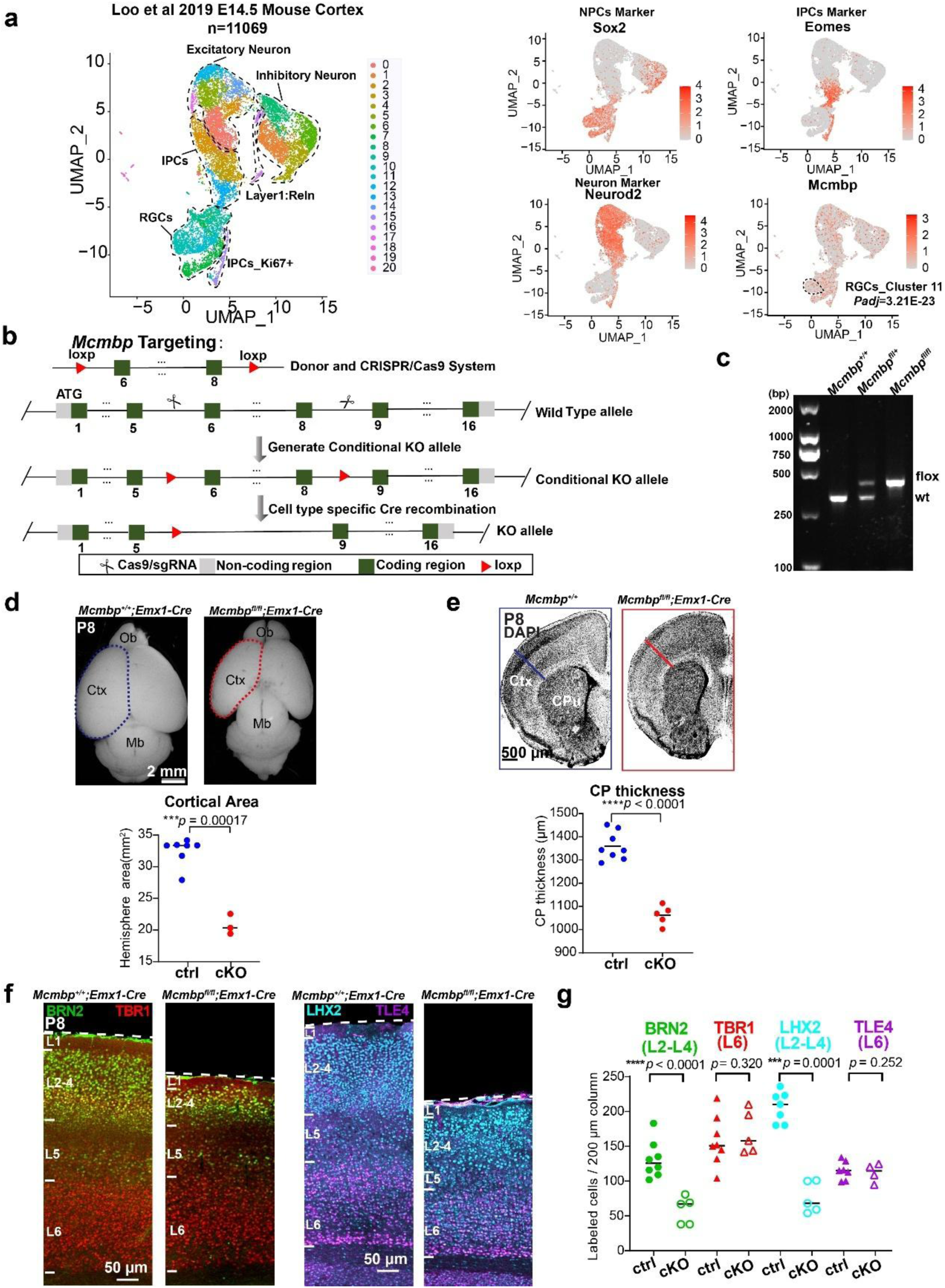
|Generation of *Mcmbp* conditional knock-out mice. **a**, Uniform manifold approximation and projection (UMAP) plot showing cell types identified in the E14.5 mouse cortex. NPC marker gene *Sox2*, IPCs mark gene *Eomes*, neuronal marker gene *Neurod2*, and *Mcmbp* expression in the identified cell types. **b**, Schematic representation of the generation of Mcmbp conditional floxed mice using CRISPR-Cas9 technology. **c**, PCR results from wild-type, heterozygous and homozygous mice. **d**, (Top panel) Dorsal view of Mcmbp^+/+^; Emx1-Cre and Mcmbpfl/fl; Emx1-Cre (cKO) P8 brain. (Bottom panel) Cortical area was significantly reduced in cKO compared to littermate controls (ctrl). (mean, two-tailed unpaired t-test, ctrl: n=7, cKO: n=3) **e**, (Top panel) DAPI staining coronal sections of Mcmbp^+/+^ and cKO P8 brain. (Bottom panel) Cortical plate thickness was significantly reduced in cKO compared to littermate controls (ctrl). (mean, two-tailed unpaired t-test, ctrl: n=8, cKO: n=5) **f**, Immunostaining of layer markers BRN2, TBR1, LHX2 and TLE4 in Mcmbp^+/+^; Emx1-Cre and cKO P8 brain. **g**, Upper layer neurons were significantly reduced in cKO compared to the controls (ctrl). (mean, two-tailed unpaired t-test, BRN2, TBR1, ctrl: n=8, cKO: n=5, LHX2, TLE4, ctrl: n=7, cKO: n=5).

**Extended Data Fig.2.**
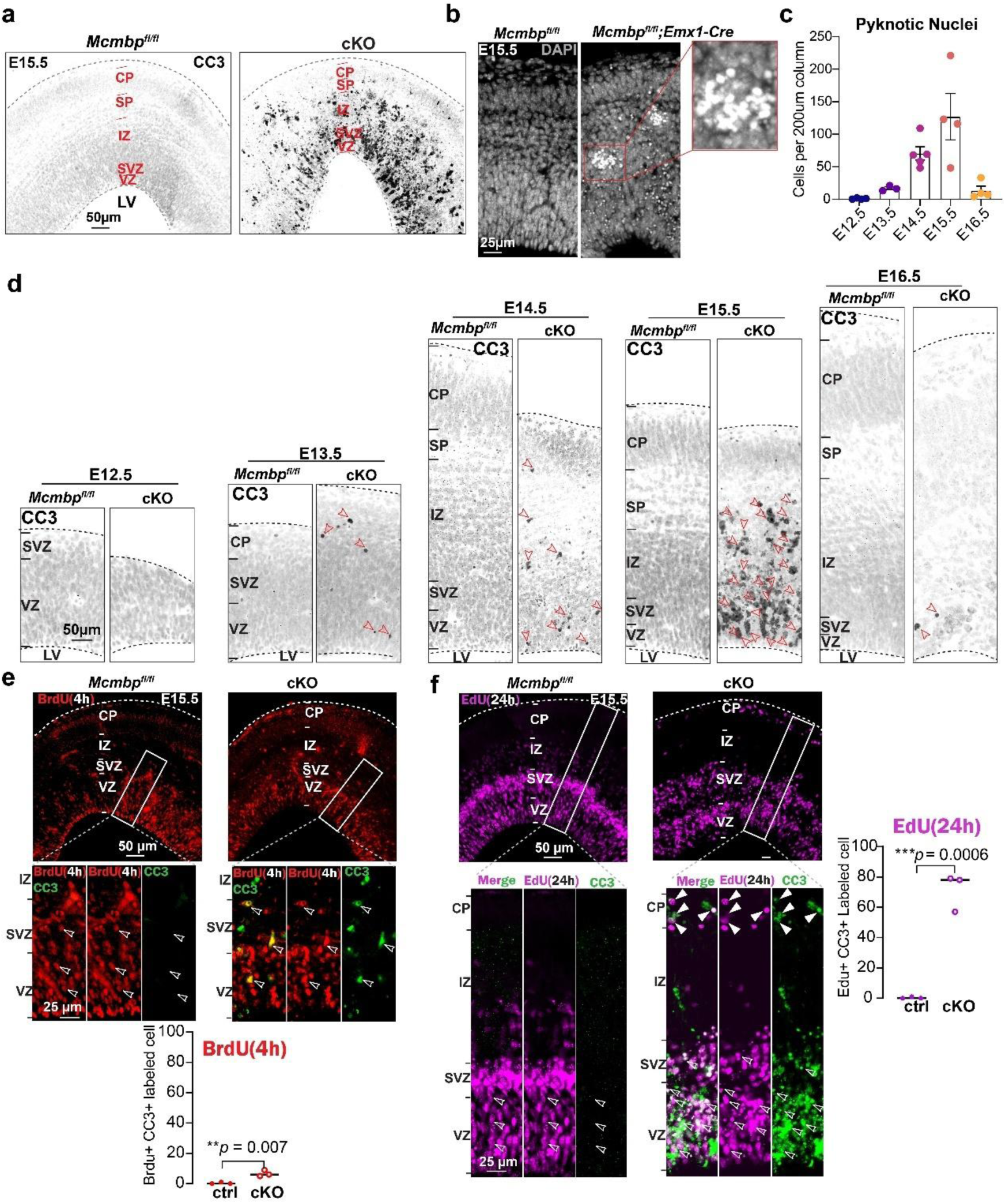
|*Mcmbp* deletion in neural progenitors leads to rapid apoptosis. **a**, CC3 immunostaining revealed extensive apoptosis in cKO at E15.5. **b**, DAPI staining revealed pyknotic nuclei in cKO cortical plate, clustered pyknotic nuclei are marked with red box. **c**, Quantification results indicated that the number of pyknotic nuclei appeared at E13.5, and reached the peak at E15.5.(E12.5: n=4, E13.5: n=3, E14.5: n=5, E15.5: n=4, E16.5: n=4) **d**, CC3 immunostaining from E12.5 to E16.5 indicated that cell apoptosis appeared at E13.5 but not at E12.5, and reached the peak at E15.5, which was coincided with the peak for neurogenesis. Red arrows indicate apoptotic cell. **e**, (Top panel) Analysis of E15.5 cortex following a 4-hour pulse of BrdU (red). CC3 staining (green) showed apoptosis in BrdU+ cells (open arrowheads) in Emx1- *Mcmbp*cKO, but not in control. (Bottom panel) Quantification results of BrdU+ and CC3+, indicated NPCs rapidly entered into the apoptosis process within 4 hours following *Mcmbp* deletion. (mean, two-tailed unpaired t-test, ctrl: n=3, cKO: n=3) **f**, (Top panel) Analysis of E15.5 cortex following a 24-hour pulse of EdU (magenta). CC3 staining (green) showed apoptosis in BrdU+ cells (open arrowheads) in Emx1- *Mcmbp*cKO, but rarely in control. (Bottom panel) Quantification results of EdU+ and CC3+, indicated massive numbers of NPCs entered into apoptosis process in cKO within 24 hours. (mean, two-tailed unpaired t-test, ctrl: n=3, cKO: n=3)

**Extended Data Fig.3.**
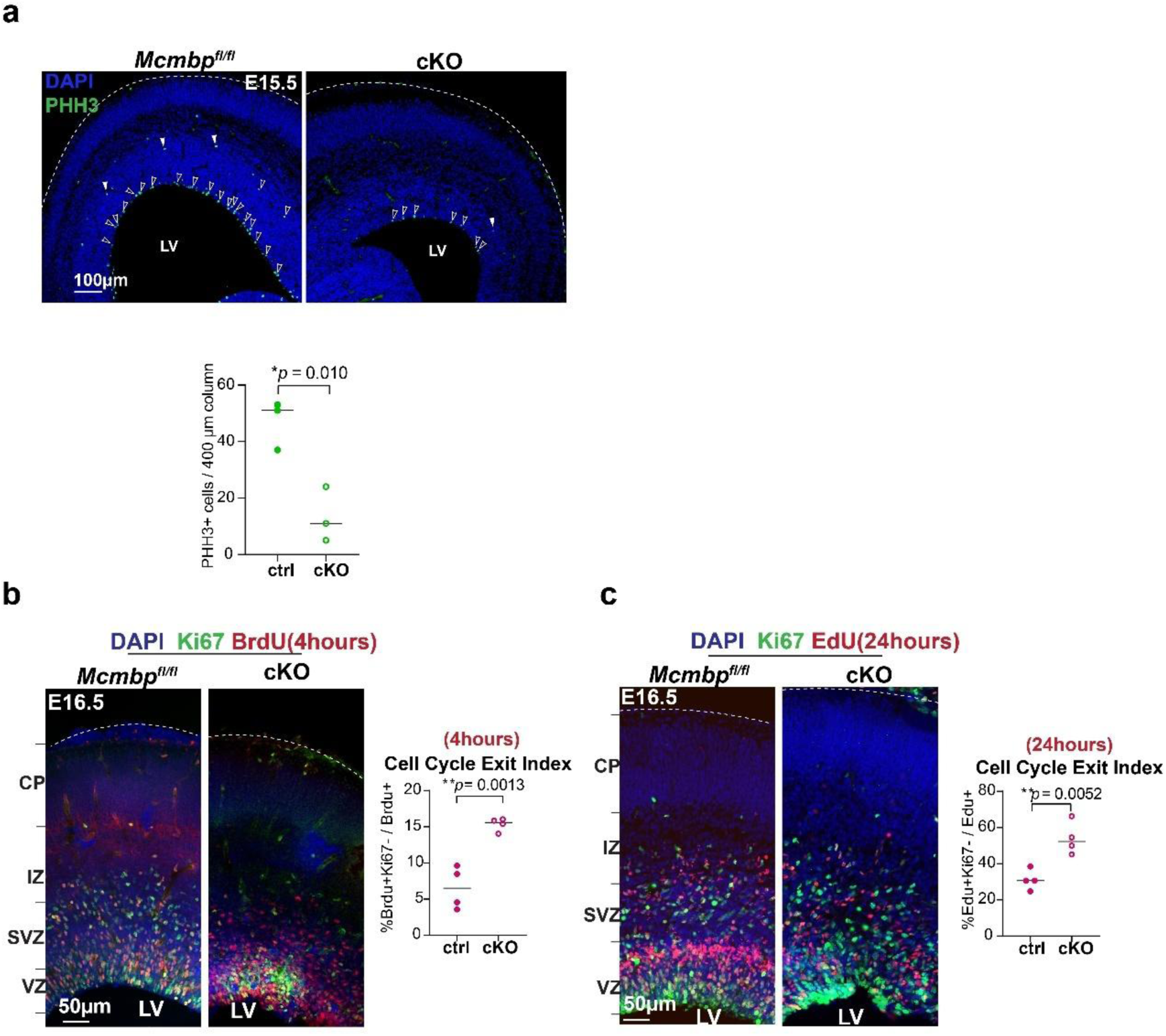
|Cell cycle analysis in neural progenitors following *Mcmbp* deletion. **a**, Representative images and a quantification of M-phase PHH3+ cells at E15.5. (mean, two-tailed unpaired t-test, ctrl: n=3, cKO: n=3) **b**, Representative images and a quantification of cell-cycle-exiting-cells following BrdU pulse for 4 hours at E15.5. (mean, two-tailed unpaired t-test, ctrl: n=4, cKO: n=4) **c**, Representative images and a quantification of cell-cycle-exiting-cells following EdU pulse for 24 hours at E15.5. (mean, two-tailed unpaired t-test, ctrl: n=4, cKO: n=4)

**Extended Data Fig.4.**
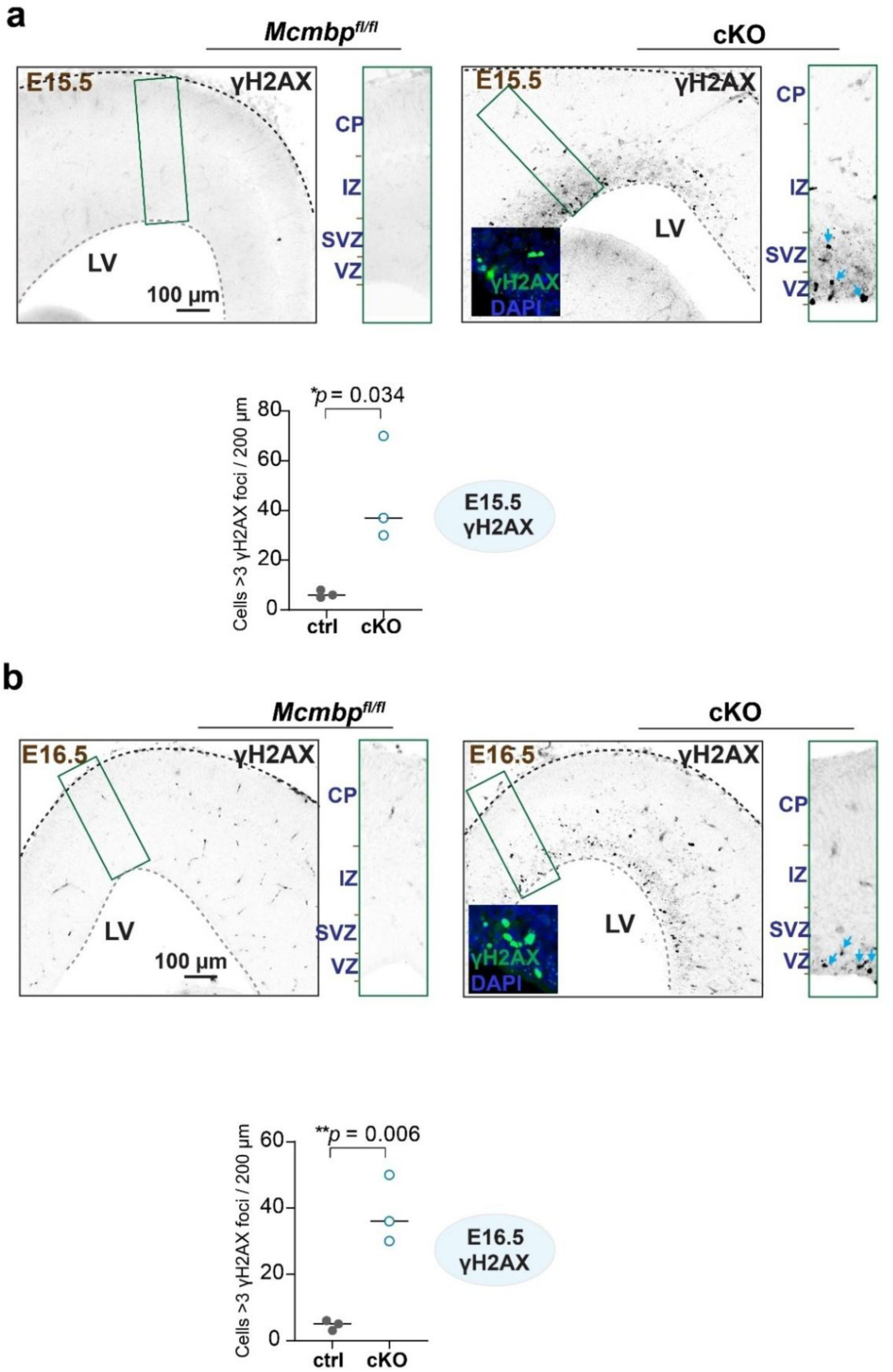
|DNA damage analysis in *Mcmbp* cKO. **a**, Representative images and quantification of DNA-damaged γH2AX+ cells at E15.5. (mean, two-tailed unpaired t-test, ctrl: n=3, cKO: n=3). **b**, Representative images and quantification of DNA-damaged γH2AX+ cells at E16.5. (mean, two-tailed unpaired t-test, ctrl: n=3, cKO: n=3).

**Extended Data Fig.5.**
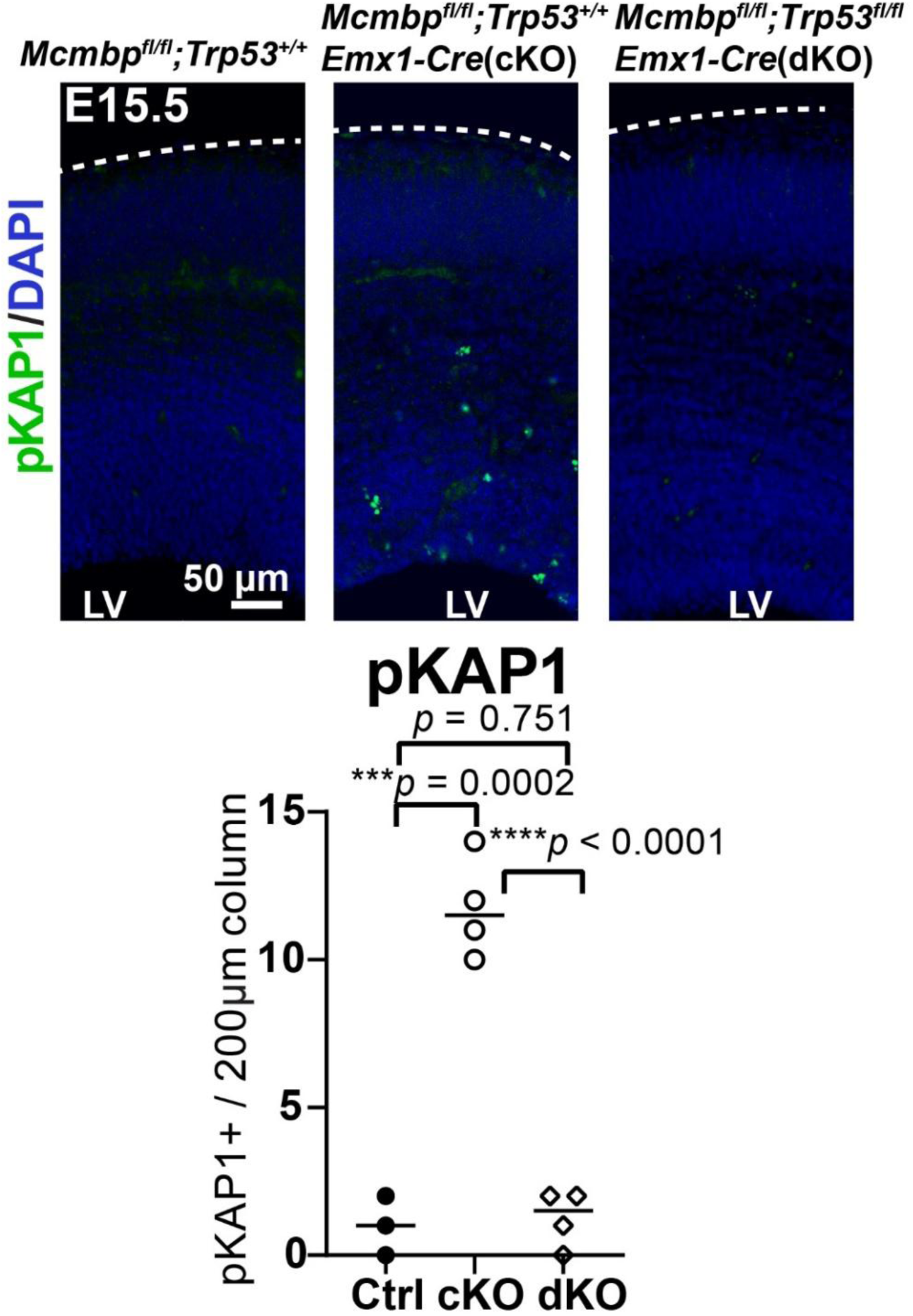
|Representative images and quantification of pKAP1+ cells at E15.5. (mean, two-tailed unpaired t-test, ctrl: n=3, cKO: n=4, dKO: n=4).

**Extended Data Fig.6.**
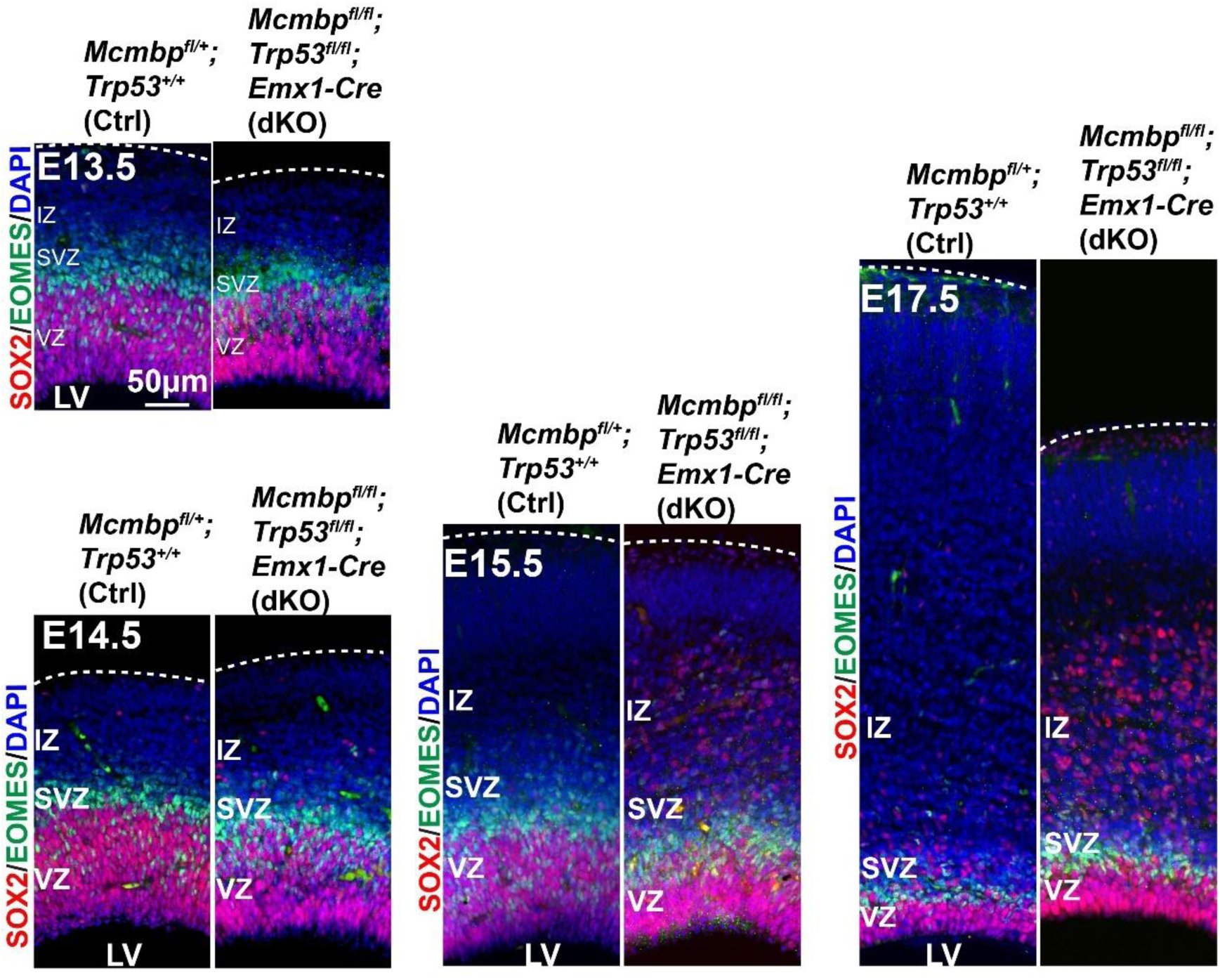
|Representative images of SOX2+ RGCs and EOMES+ IPCs in E13.5, E14.5, E15.5 and E17.5 dKO cortex.

**Extended Data Fig.7.**
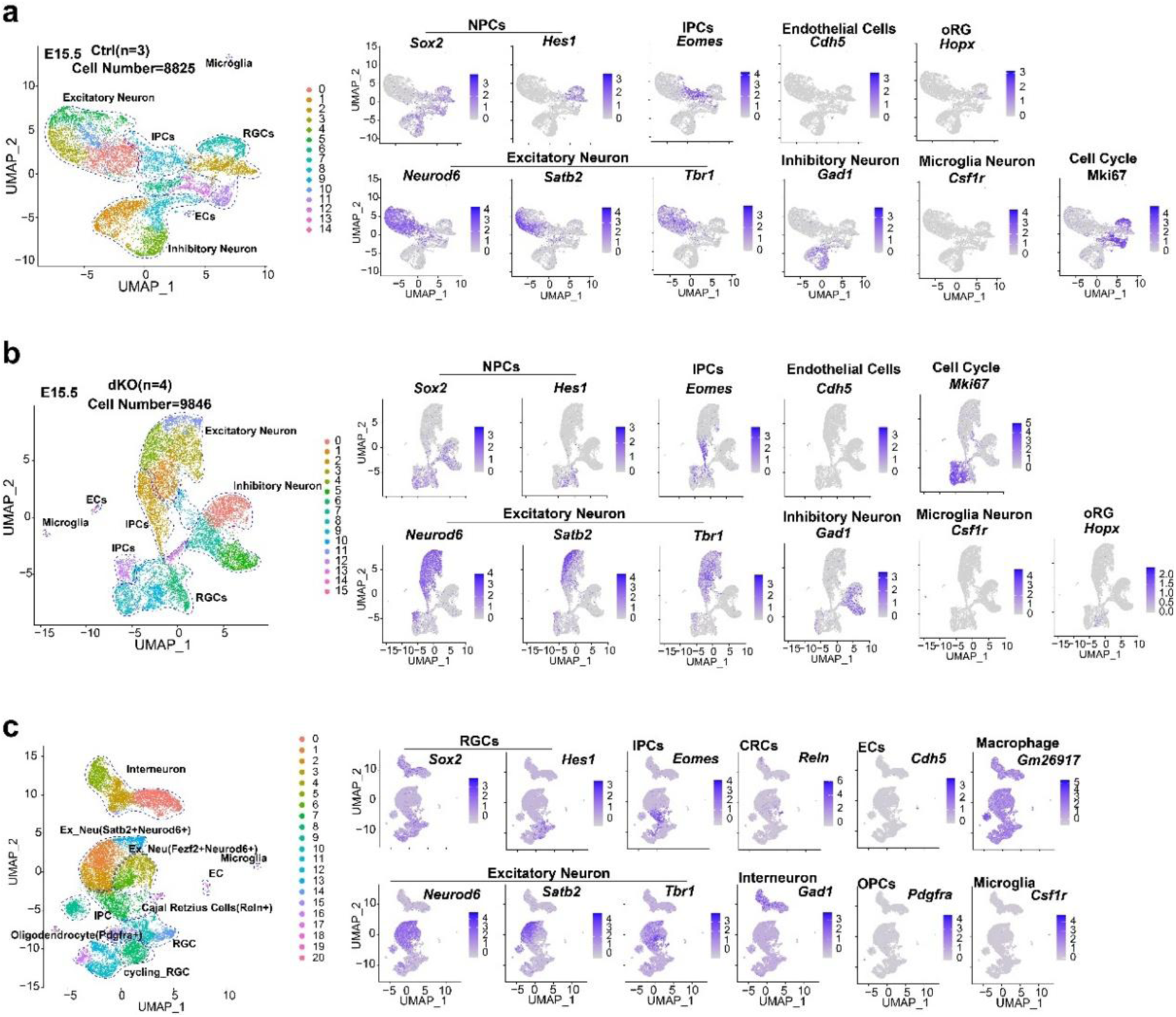
|Single-cell analysis of control and *Mcmbp/Trp53* dKO cortex. **a**, (Left panel) Uniform manifold approximation and projection (UMAP) plot showing cell types identified in the control embryonic cortex. (Right panel) Feature plot showing cell-type-specific marker genes’ expression in the controls. **b**, (Left panel) Uniform manifold approximation and projection (UMAP) plot showing cell types identified in the dKO embryonic cortex. (Right panel) Feature plot showing cell-type-specific marker genes’ expression in the controls. **c**, (Left panel) Uniform manifold approximation and projection (UMAP) plot showing cell types identified in the integrated single-cell data from control and dKO cortex. (Right panel) Feature plots of RGCs, IPCs, CRCs, ECs, macrophages, excitatory neurons, interneurons, OPCs, and microglia markers in the integrated single-cell data.

**Extended Data Fig.8.**
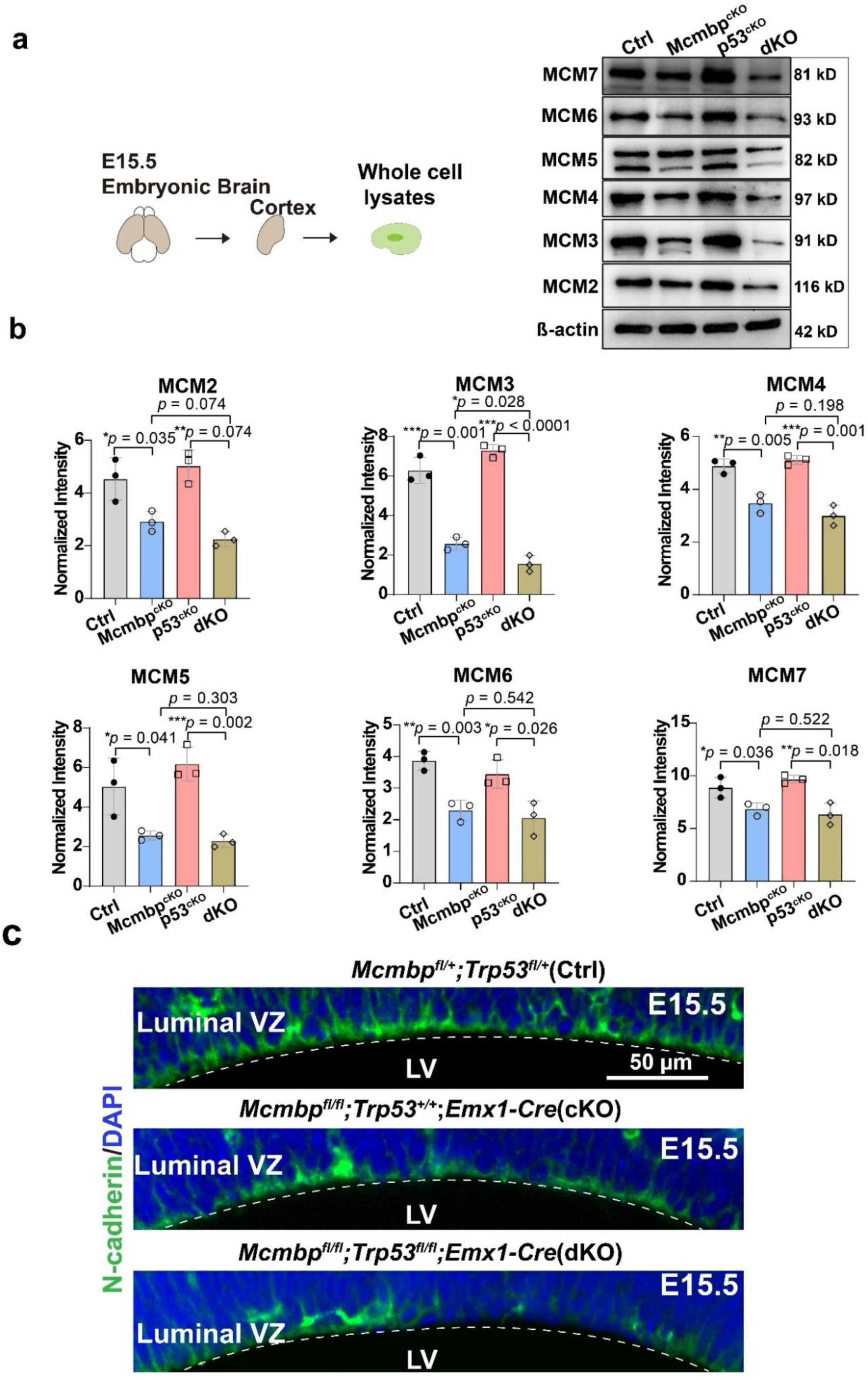
|MCM complex subunit expression and cell adhesion analysis in *Mcmbp/Trp53* dKO. **a-b**, Western blotting of whole-cell lysates and quantitative analysis identifying MCM3 as the only MCM subunit significantly repressed in dKO. (mean, two-tailed unpaired t- test, ctrl: n=3, Mcmbp-cKO: n=3, p53-cKO: n=3, dKO: n=3) **c**, Representative images of adhesion marker N-cadherin+ at E15.5 dKO cortex.

**Extended Data Fig.9.**
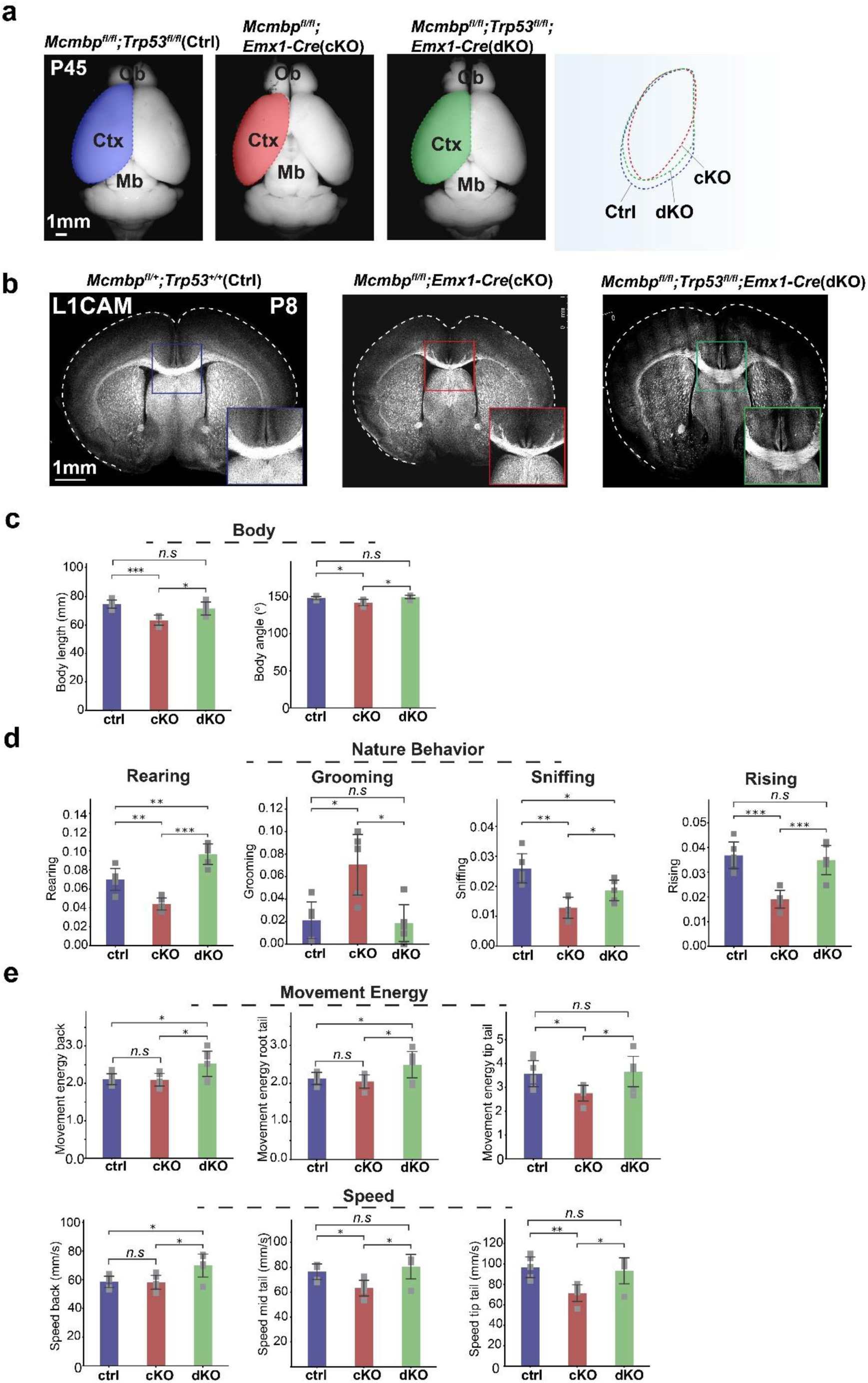
|Fork speeding acceleration leads to an anxiety-like behavior in *Mcmbp* cKO and *Mcmbp/Trp53* dKO. **a**, Images of control, cKO, and dKO brains at P45. **b**, Immunostaining of L1 cam to show corpus callosum in P8 control, cKO, and dKO. c, Bar plots of body length and angle in controls, cKO, and dKO. (Welch’s t-test, ctrl, cKO, dKO: n=5). **d**, Natural behavior analysis in controls, cKO, and dKO. (Welch’s t-test, ctrl, cKO, dKO: n=5). **e**, Bar plots of movement energy and speed in controls, cKO, and dKO. (Welch’s t-test, ctrl, cKO, dKO: n=5).

